# A moving target: non-stationary selection governs unsupervised prediction of viral fitness

**DOI:** 10.64898/2026.09.17.752359

**Authors:** Stéphane Aris-Brosou, Matthieu Vilain

## Abstract

Anticipating how mutations change viral fitness is central to genomic surveillance and vaccine design. However, the phenotypic data behind the most accurate supervised variant-effect predictors is unavailable for most emerging pathogens. It is therefore unclear how a label-free, unsupervised approach would perform using only sequences, their evolutionary history, and their protein structure. To address this outstanding question, we assemble a modular, fully unsupervised pipeline that estimates key interpretable terms: intrinsic replicative fitness, antigenic escape, and realized growth. Each term is produced by more than one estimator, so that the fitness predictor becomes a testable model choice. Benchmarking the intrinsic term on 21 viral deep-mutational-scanning assays from ProteinGym, we find that a 650-million-parameter single-sequence protein language model (PLM) is a weak predictor of viral fitness, whereas a simple site-independent alignment model more than doubles performance, with the largest gains obtained for the antigenic surface proteins not included in the training of the PLM. On the other hand, PMLs based on protein alignment (MSA Transformer, Tranception) regain the accuracy lost with the single-sequence PML, but do not significantly exceed the simple alignment. Therefore, the most important feature to predict viral fitness is the protein alignment, not model scale or architecture. Our central result is evolutionary: using dated sequences of SARS-CoV-2 spike and influenza H3N2 hemagglutinin, we find that the epoch of the alignment is itself a virus-specific determinant of accuracy. This demonstrates that selection is a non-stationary process, in a manner that is itself virus-specific. A phylogenetic mutation-selection estimator of fitness does not match the far simpler and less computationally expensive alignment model. Altogether, this work shows that unsupervised viral fitness prediction is as much an evolutionary problem as a modeling one.

## Introduction

Predicting the phenotypic consequences of amino-acid substitutions is a long-standing goal of molecular evolution, and an increasingly practical one for evolving viruses. Indeed, anticipating which mutations increase replicative fitness or confer antigenic escape (the ability to evade existing immunity) shall provide us with guidelines on surveillance, therapeutic design, and vaccine-strain selection. However, the most accurate variant-effect predictors are supervised: they learn from deep mutational scanning (DMS) or antibody-escape experiments that measure thousands of substitutions directly [1–3]. While such data are powerful, they are scarce, and rarely available for emerging pathogens when predictions are direly needed. This motivates unsupervised scoring, that is, estimating the effect of a mutation from sequence, evolutionary history, and structure alone, with no phenotypic labels. In recent years, protein language models (PLMs) have become the most prominent tools for fitness prediction, as they predict variant effects from sequence data alone.

However, this expectation for viral proteins faces two limitations. First, while trained on hundreds of millions of natural sequences, PLMs assign each substitution a zero-shot likelihood, that is, a score produced with no protein-specific fitting or labeled examples [4–6]. As a result, these approaches perform very well when benchmarked on large, mostly cellular, sets of variant effects [7–10]. But, these results do not obviously carry over to fast-evolving viral surface proteins, such as SARS-CoV-2 spike or the influenza hemagglutinin, precisely because such proteins are sparsely represented in these models’ training sets. The second limitation may be more subtle: viral selection is not stationary [11]. The sequences available to any unsupervised method are only a snapshot of an evolutionary process still in motion, so when those sequences were collected may affect how well viral fitness can be predicted. We show here that both limitations are critical, with the latter, the non-stationarity, ending up being a critical determinant of accuracy, one that static benchmarks are structurally unable to reveal.

These attempts at predicting fitness are not, however, without precedent. Estimating intrinsic (replicative) fitness, that is, how well a protein performs its function, from sequence data alone has a long history. A rich body of unsupervised methods estimate fitness from the constraints purifying selection leaves in a protein family: direct-coupling models of coevolution [12], variational autoencoders over alignments [13, 14], as well as from global epistatic models that capture non-additive interactions among mutations [15]. PLMs extend the possibility of inferring constraints from a protein’s evolutionary relatives by learning those constraints across the entire protein universe [4, 6]. In contrast, alignment-based models (such as the MSA Transformer [16] and Tranception [17]) take an explicit protein alignment as input when scoring a mutation, rather than relying on a single sequence during training. Systematic benchmarks that compare these predictors at scale typically rank these alignment-based models near the top, though they do so largely on human and microbial proteins [7, 9, 10].

All of these methods capture only one aspect of viral fitness: intrinsic, replicative fitness. A second aspect matters just as much for an immune-evading virus: antigenic escape, i.e. the ability to evade host antibodies, which is itself a component of fitness, as these escape mutations often carry a replicative cost [18]. However, purifying selection on its own cannot capture this aspect, therefore complementary features need to be inferred in one of two ways: either by combining an intrinsic-fitness term with structural and chemical information to identify disruptive substitutions, as in EVEscape [19]; or by inferring antigenic escape directly from sequence with language models [20]. These inference methods build on a large body of experimental escape mappings for SARS-CoV-2 [1, 3] and influenza [2], and on the broader description of how spike mutations drive immune escape and variant fitness [21]. A recurring conclusion in this body of work is that the constraints themselves change through time: e.g., the mutational tolerance of the SARS-CoV-2 receptor-binding domain shifted measurably as the virus evolved [11].

Such selection shifts are, in molecular evolution, not a novel observation. Site-specific substitution processes change across a tree, a phenomenon termed heterotachy [22], and both amino-acid composition and the exchange process can be non-homogeneous and non-stationary [23–25]. Models that let a site switch between variable and invariable states (the covarion models) [26, 27], that mix amino-acid profiles across sites [28], or that let those profiles themselves change through time [29] were built precisely to accommodate this behavior. What has not been investigated though, is the consequences of such non-stationarity on unsupervised fitness prediction models, which all assume that the alignment behind a score is, implicitly, constant in time.

To address this outstanding knowledge gap, we resort to an empirical approach. Instead of using a single predictor, we combine multiple unsupervised scores or terms: an intrinsic (replicative) fitness term, an antigenic-escape term, and a realized-growth term. We then let each component be produced by one of several unsupervised estimators, so that the choice of estimator becomes something we can test rather than assume. Practically, we score each candidate mutation for how well it preserves the protein’s function (the intrinsic term), how likely it is to help the virus evade antibodies (the antigenic term), and how fast the variants carrying it actually spread (the growth term). We describe the full pipeline below and keep it distinct from the one component we put to the test here: the intrinsic term. It is the only one we can check without new experiments, because the DMS assays introduced above already measure the mutational fitness effects an intrinsic-fitness score aims to predict. We evaluate this approach with three unsupervised estimators: the zero-shot likelihood of a single-sequence PLM (ESM-2), a site-independent alignment position-specific scoring matrix (PSSM; a tally of the amino acids seen at each alignment column), and a Bayesian phylogenetic mutation-selection model (which infers each site’s amino-acid preferences from the pattern of substitutions along the evolutionary tree). Benchmarking on the viral subset of ProteinGym [7] reveals the first limitation highlighted above. Using dated, same-virus samples of SARS-CoV-2 spike and influenza H3N2 hemagglutinin then exposes the inhomogeneity of the evolutionary process, linking the accuracy of an unsupervised score to the non-stationarity of the underlying substitution process [22, 25]. That single-sequence models do worse than alignment-based methods on shallow or unusual protein families is already known: large benchmarks report the same once performance is broken down by alignment depth or taxon [7]. Our contribution is different: for viral proteins, the actual sequences included in an alignment and their collection dates can matter more than the model used to score them.

## Results

Our strategy follows from the limitations above. Because a label-free score is only as good as its inputs, we decompose fitness into a few interpretable terms and treat each estimator as a testable modeling choice, not an assumption (see S1 Fig for an overview). Here, we benchmark only the intrinsic replicative fitness term, since it is the only one with experimental DMS validation. We evaluate it with three interchangeable estimators: ESM-2 zero-shot likelihood, a site-independent PSSM, and a Bayesian phylogenetic mutation-selection model. The remaining framework components, including structure-based accessibility, biochemical dissimilarity, additive and multiplicative fusion, sign-of-selection calibration, and surveillance growth, are defined in Materials and methods but are not scored against real phenotypes in this work; they are tested only on simulated data (S1 Text) and left for future validation. The empirical benchmark uses the pre-trained ESM-2 (650M) weights on 21 viral ProteinGym assays [7] and one non-viral control (Table 1, Fig 1).

**Table 1:** Empirical zero-shot benchmark on ProteinGym. Spearman correlation against measured DMS scores for the single-sequence language model (ESM-2, 650M) and the alignment-based site-independent model (PSSM). Viral aggregates are over 21 assays scored with our own harness. The non-viral aggregate is over the 186 non-viral ProteinGym assays (human, prokaryotic, eukaryotic) using ProteinGym’s released ESM-2 (650M) and site-independent scores, which match our harness to within 0.02 (Table 2); TEM-1 *β*-lactamase is one such assay, scored with our harness. Per-assay viral values are shown in Fig 1.

| Assay set | $n$ | ESM-2 | PSSM |
| --- | --- | --- | --- |
| Viral assays (mean) | 21 | 0.15 | 0.39 |
| Viral assays (median) | 21 | 0.08 | 0.38 |
| HIV envelope (Haddox_2018, BF520) | 1 | 0.08 | 0.50 |
| SARS-CoV-2 spike binding | 1 | −0.02 | 0.13 |
| Influenza HA (Doud_2016) | 1 | 0.51 | 0.50 |
| Non-viral assays (mean) | 186 | 0.47 | 0.37 |
| Non-viral control (BLAT_ECOLX_Firnberg_2014) | 1 | 0.74 | 0.49 |

**Figure 1:**
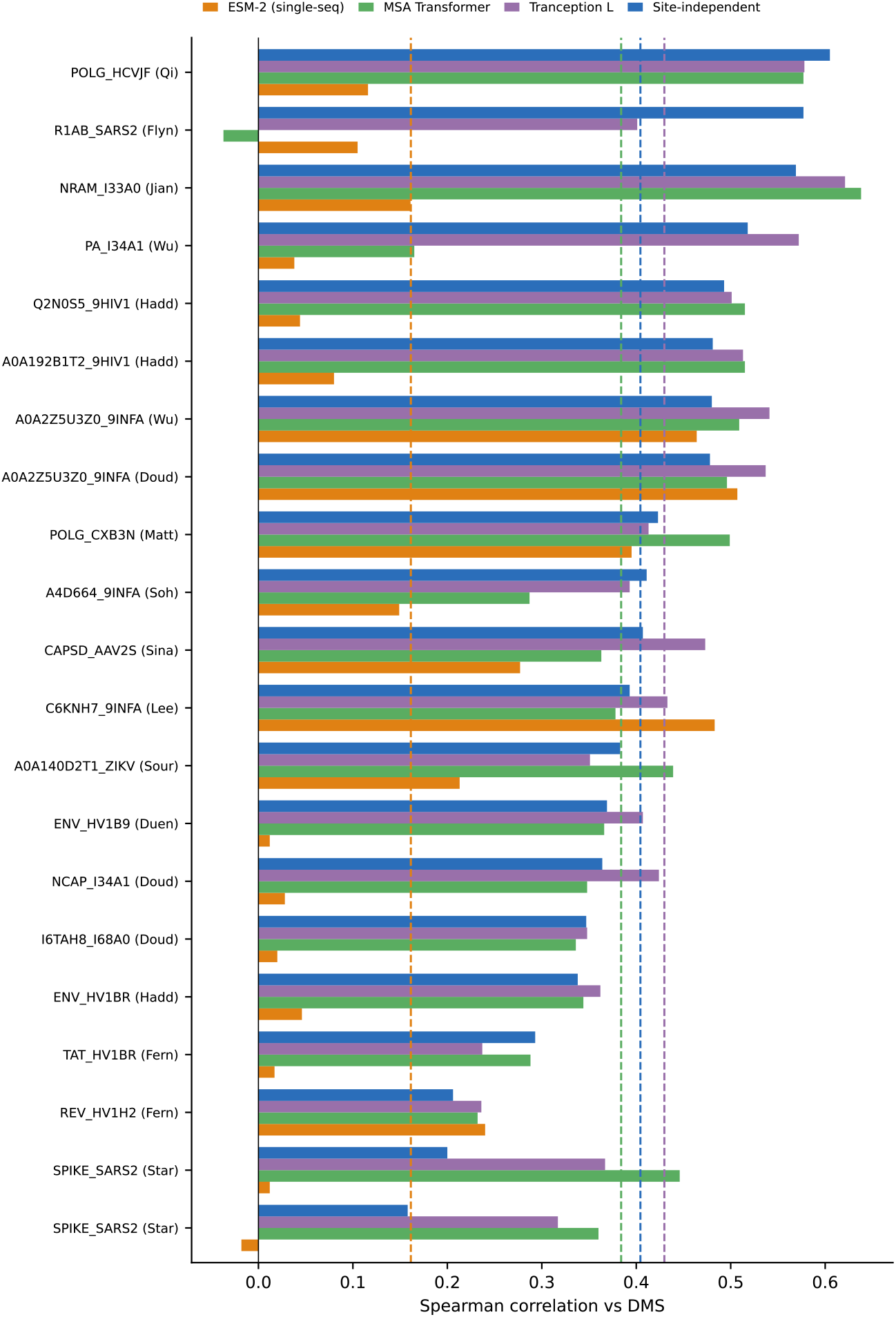
Single-sequence versus alignment-based scoring across the viral assays. Per-assay Spearman correlation against experimental DMS for four zero-shot models on the 21 ProteinGym viral assays: the single-sequence language model ESM-2 (650M), the alignment-conditioned MSA Transformer, the retrieval-augmented Tranception, and a site-independent alignment baseline. Values are ProteinGym’s released per-assay scores; dashed lines mark the per-model means (ESM-2 0.16; MSA Transformer 0.38; Tranception 0.43; site-independent 0.40). The single-sequence model is lowest on nearly every assay, whereas the two alignment-conditioned language models track the simple alignment baseline: the decisive ingredient is the protein alignment, not model scale.

If alignment is more informative than the sequence model, the key question is which alignment to use. To address this, we assemble dated samples and make evolutionary change explicit by building the intrinsic term from sequences in successive collection-date windows, then measuring how well each predicts a fixed assay. We track changes in predictive power over time and test whether a phylogenetic estimator that separates selection from neutral mutation performs better than a simple alignment count.

### Empirical benchmark on ProteinGym

To measure the predictive accuracy of the intrinsic-fitness term with a real language model, we scored ProteinGym substitution assays with ESM-2 (650M parameters) masked-marginal likelihoods (the log-likelihood ratio of the mutant to the wild-type residue in the same sequence context; Eq 1, Materials and methods) in intrinsic-fitness mode. We then computed the per-assay Spearman rank correlation against the measured DMS scores. Note that this is a rank-based measure of agreement, that scores +1 for a perfect monotonic match between predicted fitness and measured DMS scores, and 0 for none. As a positive control, a non-viral enzyme assay (TEM-1 *β*-lactamase, `BLAT ECOLX Firnberg 2014`) gave a Spearman correlation of 0.74, which is in line with published ESM-2 performance. This shows that our approach behaves as expected on a protein benchmarked in the original model.

Next, since estimating viral fitness is our prime objective, we examined 21 viral assays Table S1. Nineteen of the 21 assays measure viral replication, and only the two SARS-CoV-2 spike RBD assays measure binding to the host receptor ACE2, and surface expression. Here on viral data, the single-sequence language-model term is both weaker and highly heterogeneous (Table 1, Fig 1): the mean Spearman correlation is only 0.15 (median 0.08), ranging from 0.51 for an influenza hemagglutinin replication assay down to *−* 0.02 for the SARS-CoV-2 spike receptor-binding-domain (RBD) binding assay. Performance is highest for replication assays of conserved proteins, but collapses toward zero for antigenic and receptor-binding phenotypes of fast-evolving surface proteins (SARS-CoV-2 spike binding and expression, HIV envelope, HIV Tat, all ≤0.08). On the spike RBD binding assay, even a trivial substitution-matrix baseline (a fixed BLOSUM62 exchangeability score that uses no protein or alignment information, distinct from the alignment PSSM introduced next; *ρ* = 0.31) outperforms ESM-2 (*ρ* = *−* 0.02; Fig 2A).

**Figure 2:**
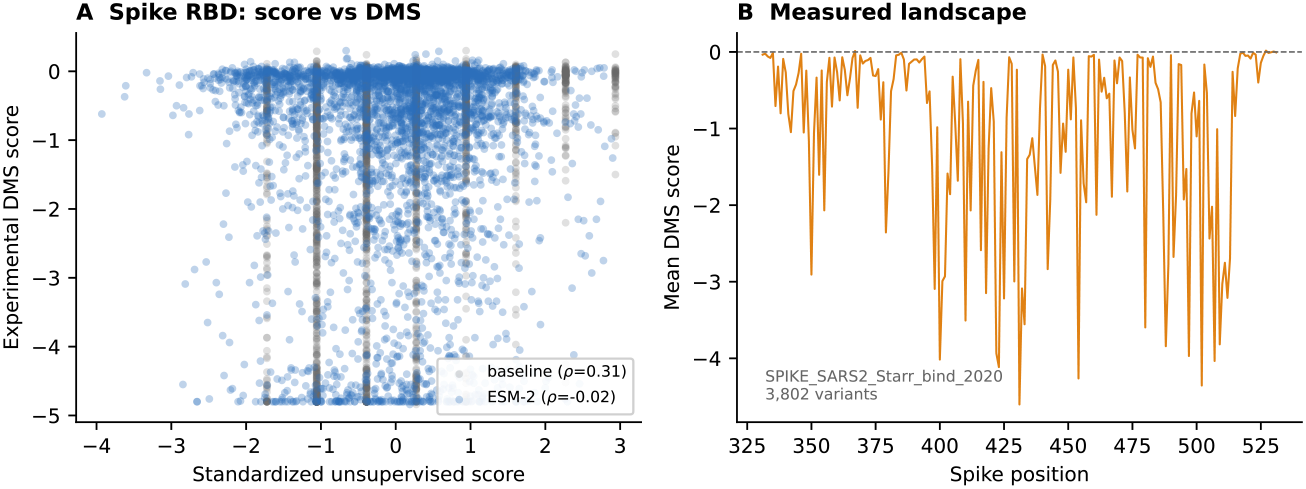
Real-data benchmark on the SARS-CoV-2 spike RBD. (A) Unsupervised score against the experimental deep mutational scanning score for every measured variant of the ProteinGym `SPIKE SARS2 Starr bind 2020` assay, on a standardized score axis: the ESM-2 masked-marginal score (*ρ* = *−* 0.02) and, for comparison, the substitution-matrix baseline (*ρ* = 0.31). The simple baseline outperforms the language model on this antigenic assay. (B) The measured mutational landscape (mean DMS score per residue) across the assayed region.

Next, we replaced the single-sequence term with the alignment-based site-independent model (PSSM), scored again from the ProteinGym multiple-sequence alignments under a single, fixed rule applied to every assay: up to the first 20,000 aligned sequences, with no per-sequence reweighting, no identity-based selection, and a pseudocount of 0.5 (a small constant added to every residue count to avoid zeros), and never tuned per assay against the DMS. Performance rises sharply across the same assays: the mean Spearman correlation goes from 0.15 to 0.39 (median 0.08 to 0.38), and the alignment-based model wins on 17 of 21 assays, with a mean improvement of +0.24 (Table 1). The advantage is significant across assays by a one-sided paired Wilcoxon signed-rank test (*p* = 8 ×10^*−*5^; sign test *p* = 3.6 ×10^*−*3^, *n* = 21). Because two assays probe the same protein, we also tested at the protein level: across the 19 distinct proteins, the advantage is essentially unchanged (mean PSSM 0.40 vs ESM-2 0.15; PSSM higher for 16 of 19 proteins; paired Wilcoxon *p* = 1.3 ×10^*−*4^). This shows that these results are not an artifact of repeated assays of the same protein.

The gains from replacing ESM-2 with the PSSM are largest precisely for the antigenic and surface proteins on which the language model failed. The observed gains from ESM-2 to PSSM are for the Haddox 2018 HIV envelope assays (0.04-0.08 →0.50-0.52), influenza neuraminidase and Polymerase Acidic proteins (PA; →0.58-0.62), and SARS-CoV-2 spike (≤0.02 →0.13-0.29). The two models are comparable on the conserved influenza hemagglutinin assays where the language model already did well.

Now, on non-viral proteins, the ordering reverses. Across the 186 non-viral substitution assays in ProteinGym (human, prokaryotic, and eukaryotic), single-sequence ESM-2 (650M) significantly exceeds the site-independent alignment model (mean Spearman 0.47 vs 0.37; ESM-2 higher on 152 of 186 assays; paired Wilcoxon *p* = 2 ×10^−15^; Table 1). This is the opposite of the viral result, and the TEM-1 *β*-lactamase control above (0.74 vs 0.49) is one such case. These non-viral values come from ProteinGym’s released per-assay scores. On the 21 viral assays, those released scores reproduce our own ESM-2 and site-independent numbers to within 0.02 units of correlation (Table 2 reports that viral cross-check). Therefore, the released numbers and those presented here are directly comparable. The advantage of the alignment model is hence specific to viral proteins rather than general. All viral PSSM mappings had high identity between the alignment’s focus (reference) sequence and the assay’s target sequence (focus-to-target identity ≥ 0.998), and high coverage of the assayed positions by alignment columns (per-assay coverage ≥ 0.947). This shows that the comparison is not confounded by alignment artifacts (per-assay values in Table S2).

**Table 2:** Alignment-conditioned language models versus the site-independent baseline on the 21 viral assays. Mean and median Spearman correlation with the experimental DMS, from ProteinGym’s official released per-assay scores (one internally consistent source). Single-sequence ESM-2 is far below the others (paired Wilcoxon *p<* 10 ^*−* 3^ vs the site-independent model); the MSA Transformer is indistinguishable from the site-independent model (*p* = 0.59) and Tranception only marginally above it (*p* = 0.04).

| Model | Mean $\rho$ | Median $\rho$ |
| --- | --- | --- |
| ESM-2 650M (single-sequence) | 0.16 | 0.11 |
| MSA Transformer | 0.38 | 0.37 |
| Tranception L (retrieval) | 0.43 | 0.41 |
| Site-independent (alignment) | 0.40 | 0.41 |

We emphasize that ProteinGym only probes the intrinsic term: these assays measure replication, binding, or expression fitness, not antibody escape or epidemiological growth, which require the escape-mapping and surveillance data used to validate the other components. The benchmark scores the intrinsic term on its own, and applies no term fusion, so that the additive and multiplicative fusion modes are not exercised here. Contrasting the two modes would require escape- or growth-labeled data, which is outside the scope of the present work.

To gauge if this weakness is a property of language models in general, or more specifically of the scoring from a single sequence, we compared these results against two alignment-conditioned language models that, unlike ESM-2, are given a protein alignment at inference: the MSA Transformer [16] and the retrieval-augmented Tranception [17]. Using their official per-assay scores released with ProteinGym on the same 21 viral assays (Table 2, Fig 1), our independent ESM-2 run reproduces ProteinGym’s to within 0.01 units of correlation (mean 0.154 vs 0.161), and our PSSM matches their site-independent baseline to within 0.02 units. These comparable numbers show that both alignment-conditioned language models recover nearly all of the accuracy that the single-sequence model lacks: the MSA Transformer reaches a mean Spearman of 0.38, statistically indistinguishable from the simple site-independent alignment model (0.40; paired Wilcoxon *p* = 0.59). Tranception reaches 0.43, only marginally above it (+0.025, paired mean difference; *p* = 0.04), versus 0.16 for single-sequence ESM-2 (*p<* 10 ^*−* 3^ below the alignment baseline). A single-sequence PLM falls well below the alignment-conditioned PLMs, which only restore performance to roughly that of a site-independent count. However, none of these models clearly exceeds that simple baseline on viral proteins (Fig 3).

**Figure 3:**
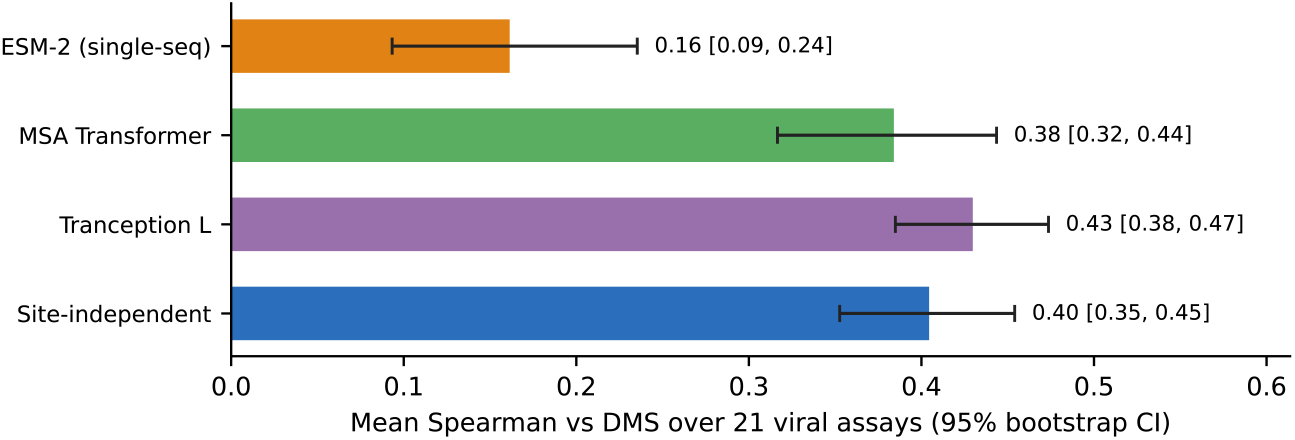
Alignment is the decisive ingredient, not model scale. Mean Spearman correlation with the experimental DMS over the 21 viral assays, with 95% bootstrap confidence intervals (10^4^ resamples of the assays), from ProteinGym’s released per-assay scores. The single-sequence ESM-2 model sits well below the others with a non-overlapping interval, whereas the two alignment-conditioned language models (MSA Transformer, Tranception) and the simple site-independent alignment model overlap.

### Prediction quality depends on the collection date

An important caveat with what was tested so far is that the benchmark alignments above are based on undated homologs, which cannot reveal any time structure in the protein family. We therefore assembled dated samples (2,800 SARS-CoV-2 spike and 2,730 H3N2 hemagglutinin sequences carrying collection dates). For a series of collection-date windows (cumulative, and sliding *±*6 months) that we call epochs, we then built a PSSM from only the sequences within each window, and scored it against the same fixed DMS data (Materials and methods). We found that prediction depends strongly on the collection epoch in both viruses, but the two proteins behave in contrasting ways (Fig 4). The two phenotypes also differ: the spike assay measures ACE2-binding affinity of the receptor-binding domain, whereas the hemagglutinin assay measures viral replication, a more global constraint on protein function.

**Figure 4:**
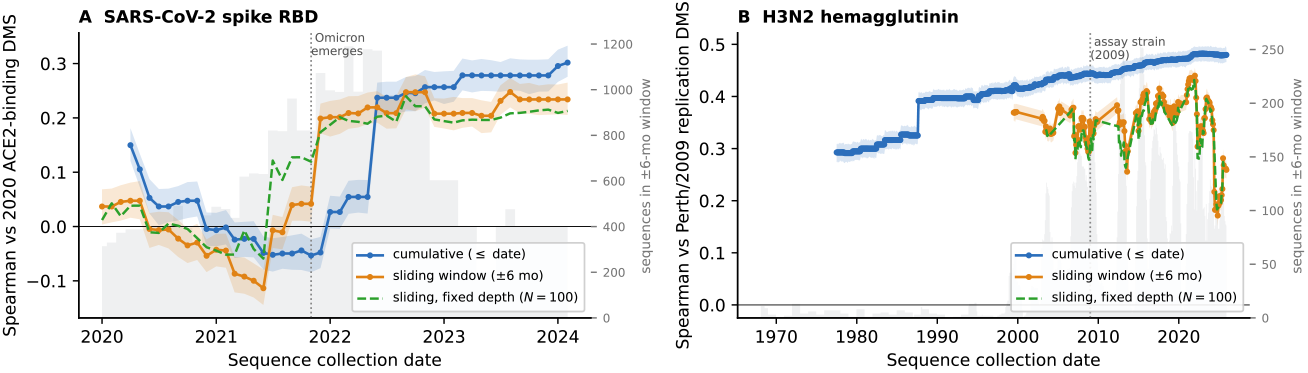
Prediction of a fixed DMS depends on the collection date of the alignment, in a virus-specific way. Spearman correlation between a same-virus PSSM and a fixed deep-mutational-scanning assay, as a function of sequence collection date, for a cumulative window (sequences up to the date) and a sliding *±* 6-month window. **(A)** SARS-CoV-2 spike RBD (2,800 dated sequences) vs the Starr et al. [1] 2020 ACE2-binding assay: near zero while the RBD is invariant, then a sharp rise as Omicron diversifies the RBD (punctuated mode). **(B)** H3N2 hemagglutinin (2,730 dated sequences) vs the Lee et al. [2] Perth/2009 replication assay: positive throughout and rising gradually with accumulated diversity (continuous-drift mode). Shaded bands are 95% bootstrap confidence intervals (resampling the assayed mutations); the grey area (right axis) is the number of sequences in the sliding *±* 6-month window. The dashed green curve is a depth-matched control: every window is subsampled to a fixed *N* = 100 sequences (mean over 20 draws), holding alignment depth constant across epochs. It tracks the native sliding window closely (Spearman 0.96 for spike, 0.95 for HA), so the date dependence is not an artifact of alignments growing deeper over time.

For SARS-CoV-2 spike (Fig 4A), the time-dependence is punctuated. Through 2020 and 2021, both the cumulative and sliding-window correlations with the 2020 ACE2-binding assay sit near zero, and are transiently negative (*ρ ≈ −* 0.11 in mid-2021). This is probably because the RBD was almost invariant among circulating viruses, and the alignment therefore carried little information. Both curves rise sharply as the Omicron variants emerge in late 2021 and plateau around *ρ* = 0.20 to 0.25 through 2022 to 2024, the sliding window rising first (contemporaneous Omicron diversity), and the cumulative one following as Omicron-era sequences accumulate. The cumulative PSSM over all 2,800 dated sequences reaches *ρ* = 0.30, more than double the 0.13 obtained for the same assay from the undated cross-species homolog alignment (Table 1).

For influenza H3N2 hemagglutinin (Fig 4B), the dependence is instead continuous. Because H3N2 has circulated with continual antigenic drift since 1968, contemporaneous alignments carry mutational-tolerance signal in every epoch: the sliding-window correlation with the Perth/2009 replication assay never approaches zero, ranging over *ρ* = 0.17-0.44, and the cumulative PSSM climbs smoothly from *ρ ≈* 0.29 in the 1980s to 0.48 in the 2020s, matching the ESM-2 score for this assay (Table S2) and exceeding the homolog-alignment PSSM (0.37). The sliding window shows no peak at the 2009 assay strain (a 2005 or 2015 alignment predicts the assay about as well), indicating that the shape of the replication constraint is comparatively time-homogeneous: much the same sites are tolerated across epochs, in contrast to the epoch-specific binding constraint of the spike’s RBD. For hemagglutinin, what changes with epoch is therefore the magnitude of accuracy rather than the identity of the constrained sites, as a deeper, more diverse alignment estimates this stable constraint more precisely. In both viruses, then, the epoch of the alignment is a dominant determinant of predictive accuracy, whether because the constraint itself shifts, as for spike, or because the alignment captures more of the family’s variation around a time-invariant constraint, as for hemagglutinin, and a contemporaneous alignment matches or beats both the cross-species homolog PSSM and the language model. Bootstrap confidence intervals over the assayed mutations show that these epoch-to-epoch movements exceed sampling noise (shaded bands in Fig 4).

The pattern is not an artifact of alignment depth growing over time: a depth-matched control that subsamples every window to a fixed *N* = 100 sequences reproduces the native sliding curve almost exactly (Spearman 0.96 for spike and 0.95 for HA over the same collection-date range; Fig 4). Therefore, at constant depth, the accuracy still tracks the collection date.

### The site-specific substitution process is non-stationary

While the epoch dependence above is an indirect signature of a non-stationary process, we can also measure the inhomogeneity directly, from the dated sequences alone, without any reference to any DMS data. For each virus, we built the site-specific amino-acid frequency profile in every sliding *±* 6-month window, and computed its mean per-site Jensen-Shannon divergence (JSD), a symmetric, bounded measure of how different two frequency distributions are, from the earliest-epoch profile (Fig 5). A stationary process would yield a flat, near-zero curve. Instead, both viruses drift, in the same two modes seen in the prediction curves. For SARS-CoV-2 spike, the divergence is essentially zero throughout 2020-2021, then rises abruptly as Omicron sweeps. The earliest-versus latest-epoch profiles differ by 0.11 bits on average, with 13% of RBD sites shifting by more than 0.1 bits. This reflects a punctuated change of regime. On the other hand, for H3N2 hemagglutinin, the divergence instead climbs continuously, with a sawtooth pattern as successive antigenic clusters replace one another, to 0.08 bits (16% of sites shifted). Because these curves are computed from the alignment’s own composition, not from predictive accuracy, they establish that the underlying site-specific substitution process is itself non-stationary.

**Figure 5:**
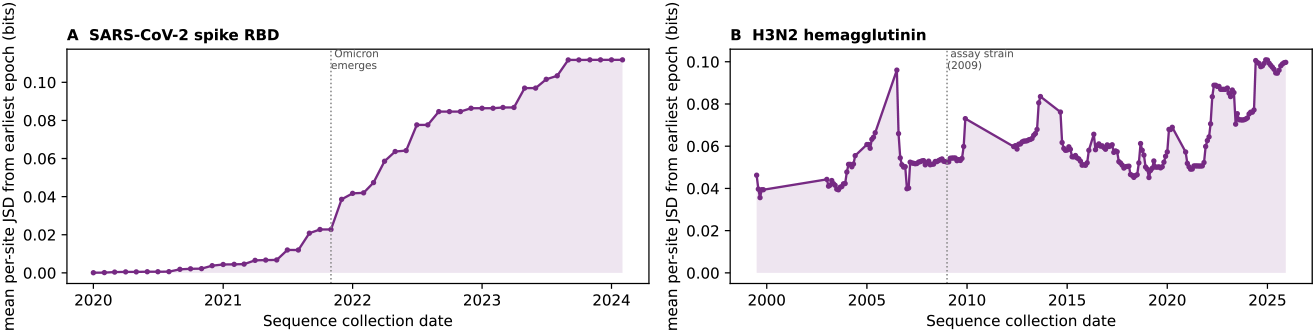
The site-specific substitution process is non-stationary, virus specifically. Mean per-site Jensen-Shannon divergence (bits) between the site-specific amino-acid frequency profile of each sliding *±* 6-month window and the earliest-epoch profile, from the dated sequences alone (no DMS). **(A)** SARS-CoV-2 spike RBD: flat near zero while the RBD is invariant, then a punctuated rise as Omicron sweeps. **(B)** H3N2 hemagglutinin: continuous drift with a sawtooth as antigenic clusters turn over.

These curves also show that the epoch-dependence of fitness prediction reflects a genuine shift in the selective regime, rather than an artifact of sampling. The curves in Fig 5 measure each window against the earliest epoch, so they track the total drift accumulated since the start of the series. On the other hand, measuring each window against the one immediately before it, which captures only the change from one epoch to the next, recovers the same two signatures: a single sharp peak at the emergence of Omicron for the SARS-CoV-2 spike, and a repeating sawtooth for H3N2 hemagglutinin (S2 Fig). Because both choices of reference give the same picture, the non-stationarity we detect is a genuine property of the sequences, not an artifact of measuring drift from the earliest epoch.

### Phylogenetic estimators do not improve on the sequence-count terms

To test whether separating selection from the neutral mutational process and correcting for shared ancestry on a tree improves prediction, we estimated the same two terms (the intrinsic and the antigenic) from a codon model, rather than from sequence counts or structure. For the intrinsic term, we fit the Rodrigue et al. [30] mutation-selection model in PhyloBayes-MPI on the codon alignment of each virus. We used 295 isolates for H3N2 hemagglutinin, and 300 for SARS-CoV-2 spike. Each dataset was analyzed on a fixed IQ-TREE topology. Note that these sample sizes are essentially limited by the cost of the site-heterogeneous Markov chain Monte Carlo (MCMC) sampler, and are deliberately far below the thousands of sequences used for the alignment and temporal analyses above. We ran two independent chains per dataset, and scored the posterior-mean site-specific fitness Δ*F* against the same DMS data. Both pairs of chains reached the sampler’s stationary regime and reproduced one another (between-chain discrepancy maxdiff *<* 0.1 across all monitored summary statistics; S3 Fig, S4 Fig). Profiles were taken as the posterior mean over both chains after the diagnostic burn-in (the initial, pre-convergence samples, which are discarded). On H3N2 HA, the tree-based fitness correlates with the Lee et al. [2] replication assay at Spearman *ρ* = 0.33 (*n* = 10,754), and on the SARS-CoV-2 spike RBD it correlates with the Starr et al. [1] ACE2-binding assay at *ρ* = 0.06 (*n* = 3,802).

To isolate the estimator from the alignment it is built on, we scored a site-independent PSSM on the same dated codon alignment and reference positions: it reaches *ρ* = 0.45 on HA and 0.21 on spike, above the mutation-selection fitness on both proteins (0.33 and 0.06) on identical data, and above the cross-species benchmark PSSM as well (0.37 and 0.13; Table S2), consistent with the value of a contemporaneous same-virus alignment. A paired bootstrap over mutations (on the shared mutation set of each assay) confirms that the gap is significant: the same-alignment PSSM exceeds the mutation-selection Δ*F* by +0.15 (95% CI [0.12, 0.19]) on the spike RBD and +0.12 ([0.10, 0.14]) on HA (both *p<* 0.001; Fig 6). On matched data, therefore, the tree-based correction for the neutral mutational supply and for shared ancestry does not improve on, and indeed falls significantly below, a simple column-frequency count.

**Figure 6:**
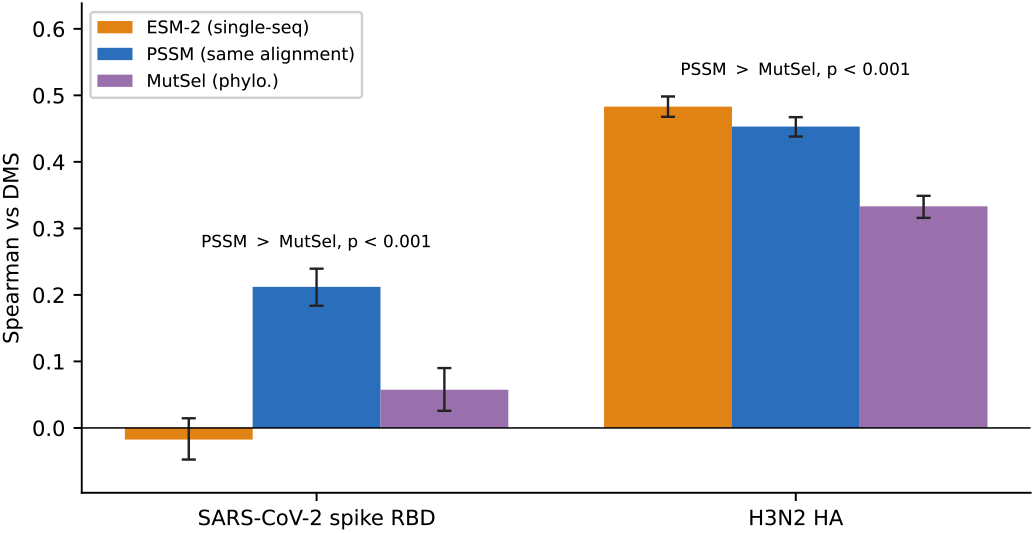
The phylogenetic mutation-selection estimator does not beat a matched alignment count. Spearman correlation with the experimental DMS for the SARS-CoV-2 spike RBD and H3N2 HA, for three estimators evaluated on the same shared mutation set of each assay: single-sequence ESM-2, a site-independent PSSM built from the same dated codon alignment used for the phylogenetic model, and the PhyloBayes mutation-selection Δ*F* . Error bars are 95% bootstrap confidence intervals (2,000 resamples of the mutations); the annotation gives the paired bootstrap test of the matched PSSM versus MutSel. On matched data the tree-based estimator falls significantly below the trivial same-alignment count for both viruses.

For the antigenic term (the escape-propensity component of the score, which up-weights immune-escape substitutions and is estimated here from a mutation-selection codon model instead of from 3D structure; Materials and methods), the four positive selection site tests (HyPhy) find small, method-dependent sets of codons under positive (diversifying) selection. On H3N2 HA the episodic-diversification test MEME [31] identifies 35 sites (FEL [32] 6, SLAC 5, FUBAR 4), consistent with the strong antigenic drift of hemagglutinin. On the more slowly diversifying spike protein, the counts are generally lower (except FUBAR) and less concordant across methods (SLAC 2, FEL 4, MEME 7, FUBAR 21). All four methods for both viruses are shown in Table S3, and the identities of the detected sites in Table S4. On the spike protein, the detected sites are enriched in the receptor-binding domain (for example positions 417, 444-446, 452, 478, and 486, all targets of neutralizing antibodies), consistent with the intended escape-site interpretation. As expected, the resulting per-site positive-selection prior is only weakly related to the experimental tolerance DMS data (Spearman ≤ 0.10 against the mean per-site DMS for both proteins). These assays score replicative or receptor-binding tolerance, not antibody escape, so a diversifying-selection signal and a tolerance measurement need not agree. The codon-model term therefore behaves as an escape-site indicator rather than a fitness predictor.

## Discussion

Scoring viral mutations without labels is not just a modeling problem: it is mainly an evolutionary one. How well a sequence-based fitness score performs depends less on the model’s sophistication than on how much evolutionary information the sequences contain and, for fast-evolving viruses, on when those sequences were sampled.

The benchmark presented here shows that performance depends strongly on the protein. Across the 186 non-viral ProteinGym assays, the language model does better than a simple site-independent alignment model, but across the 21 viral assays the alignment model performs much better. In other words, single-sequence language-model scoring is not universally reliable. It works only when the protein family is well represented in training data. When an alignment is available at inference, MSA Transformer and Tranception recover much of the accuracy that ESM-2 lacks (Table 2), showing that what matters for viral proteins is whether the scorer can access the protein family’s evolutionary variation, not which model supplies it. The ProteinGym benchmark includes 186 non-viral assays and a separate viral benchmark in which alignment-based models can outperform single-sequence ESM-2, while MSA Transformer and Tranception are alignment-aware methods. We therefore implemented a modular, fully unsupervised pipeline that combines a protein-language-model fitness term with structure- and chemistry-based escape terms akin to those of EVEscape [19]. This is a related modular framework whose fitness term derives from sequence-based evolutionary information, and whose accessibility and dissimilarity terms come from structure and amino-acid properties. Here, we evaluate only the intrinsic term against real phenotypes. The remaining components are described in Materials and methods and checked on synthetic data, but we keep those results separate.

A second result is that the right alignment to use for fitness prediction is time-dependent. By building alignments from successive collection-date windows and testing how well each predicts a fixed assay, we can measure how predictive power changes over time. This lets us gauge whether a phylogenetic estimator that separates selection from neutral mutation does better than a simple alignment count. Our main positive result is temporal. For a fast-evolving virus, the alignment behind the intrinsic term is a snapshot of a non-stationary process, so the score is tied to a particular evolutionary epoch. On dated SARS-CoV-2 spike and H3N2 hemagglutinin sequences (Fig 4), the accuracy of a same-virus PSSM on a fixed DMS changes with collection date, and the pattern reflects the virus’s evolutionary mode: punctuated for spike, where RBD-binding prediction jumps from near zero to *ρ ≈* 0.3 only after Omicron diversifies the domain; and continuous for H3N2 HA, where replication-assay prediction is positive at every timepoint and rises to *ρ ≈* 0.5 as drift accumulates. A depth-matched control shows this is not just a consequence of deeper alignments over time. Looking at the alignment composition gives the same picture: site-specific amino-acid profiles drift away from their earliest-epoch state (Fig 5), abruptly for spike at Omicron and gradually for HA, consistent with heterotachy and non-homogeneity in molecular evolution [22, 25]. Similar shifting constraints have been observed experimentally in the RBD as the virus evolved [11]. Why the two viruses differ here likely reflects what each assay measures. The spike data score ACE2-binding affinity of the receptor-binding domain [1], a function under strong directional and epistatic selection [21], so as Omicron remodeled the domain the mutations compatible with binding shifted with the sequence background, and an alignment becomes informative only once it samples that reorganized constraint. The hemagglutinin data instead score viral replication [2], a more global and conserved requirement whose mutational tolerance is comparatively stable through time, so alignments from any epoch capture it. Part of spike’s apparent epoch-dependence may also be measurement-driven, since each assay is read in a single strain background and the spike background changed far more than the hemagglutinin one over the period sampled. For viral proteins, then, where the sequences come from and when they were sampled matter as much as the model itself: a contemporaneous same-virus alignment can match or beat both the cross-species homolog PSSM and the language model.

Several limitations remain. First, although the alignment-based term more than doubles the viral correlation over the single-sequence language model, its accuracy still depends on the depth and composition of the alignment. Redundancy reweighting had little effect on these already diversity-filtered alignments (mean |Δ*ρ*| *<* 0.01), whereas the number of retained sequences had a much larger impact. On the SARS-CoV-2 spike RBD assay, correlation stayed near 0.38 up to a few thousand sequences, then dropped to 0.13 when the divergent tail of the alignment was included. In short, alignment subsampling matters more than reweighting. Restricting the alignment to sequences most similar to the assay target recovered the pathological case for spike RBD (0.13 → 0.35), but that was not consistently beneficial. In undated alignments, similarity to the target confounds evolutionary distance with oversampling redundancy, whereas dated same-virus sequences make time explicit and remove that ambiguity. Consistent with this, episodic positive selection on viral surface proteins appears to intensify through time in seasonal influenza [33] and to track variant emergence in SARS-CoV-2 [34]. This is further evidence that the selective regime behind a single alignment is not fixed in time. For spike RBD, the results are alignment-specific: the cross-species homolog PSSM gives 0.13, a depth-limited subsample 0.38, a target-similarity subset 0.35, the dated same-virus RBD alignment 0.30, and the dated codon alignment behind the phylogenetic model 0.21. Richer alignment- and retrieval-based models such as EVE, MSA Transformer, and Tranception would likely improve on the site-independent PSSM used here [7, 15]. The phylogenetic mutation-selection and codon-model positive-selection estimators are, in principle, more rigorous versions of the intrinsic and antigenic terms because they correct for mutation and shared ancestry. But on both proteins, the mutation-selection fitness estimate fell below a site-independent PSSM built from the same dated alignment (*ρ* = 0.33 vs 0.45 on H3N2 HA; 0.06 vs 0.21 on the spike RBD), so its extra computational cost is not justified for fitness prediction. One likely reason is that the fitted model assumes a single site-specific fitness profile across the tree, which our non-stationarity results in Fig 5 violate. A better remedy would be a time-heterogeneous or covarion-style substitution process [26, 27, 29], rather than a stationary one. Second, surveillance-based growth advantages are only relative, assume roughly constant advantage over the fitting window, and mix transmissibility with immune escape and epidemiological context [35, 36]. Third, structure-derived accessibility depends on having a suitable template with the correct biological assembly; when no experimental structure exists, a predicted one such as AlphaFold can be used, but that carries its own uncertainty into the accessibility term, especially for multi-chain assemblies. Fourth, the current calibration and growth estimates report point values without uncertainty, so poorly determined effects from rare variants or short windows are weighted the same as well-determined ones. The final caveat is specific to the temporal analysis. Each DMS measures a single reference strain, so correlating a date-shifted alignment against a fixed assay assumes that substitution rankings remain comparable across time. Our non-stationarity result in Fig 5 shows that this assumption is only approximate and transient: as the landscape shifts, part of the changing correlation may reflect the assay strain becoming less representative. Fully separating a changing alignment from a changing landscape would require date-matched deep mutational scans.

The essential conclusion is structural: a small set of fitness components (an intrinsic term, an antigenic term, and a realized-growth term) can each be estimated in several unsupervised ways. For viral proteins, the provenance of the sequences behind the intrinsic term matters as much as the estimator itself. Our phylogenetic tests clarify, but may not settle, the open questions. On both a continuously drifting protein (H3N2 HA) and a punctuated one (SARS-CoV-2 Spike), the mutation-selection intrinsic term did not outperform the site-independent PSSM (and actually fell below on matched data), while its codon-model antigenic counterpart identifies sites under diversifying selection that the tolerance DMS data cannot validate. The unresolved question is therefore specific: whether the codon-model antigenic prior predicts real antibody escape, which its tolerance-DMS correlation cannot test. It is also still unclear whether epoch-specific, dated alignments would systematically improve prediction for circulating viruses, as the two case studies on HA and Spike suggest. Beyond DMS, the realized-fitness components can be tested against the phylogenetic observed-vs.-expected fitness estimates of Bloom and Neher [37] and per-mutation growth advantages from surveillance [35, 36]. Methodologically, “ensembling” several estimators is likely to help [7], and propagating uncertainty in growth estimates would let calibration down-weight unreliable targets. Reconciliation of replicative constraint with antigenic novelty, the axis along which likelihood- and escape-based signals disagree [18, 20], remains the central problem for any fitness model of an immune-evading virus.

## Materials and methods

### Overview

Our goal is to score the phenotypic consequence of amino-acid substitutions in a viral protein without resorting to labeled fitness measurements. Because each fitness component can be estimated in more than one unsupervised way, we treat the estimator as a question about model choice, and compare alternative estimators against DMS data. These estimators/models are: the intrinsic fitness term from a protein language model, from an alignment position-specific scoring matrix (PSSM), or from a phylogenetic mutation-selection model; and the antigenic term from protein structure, or from a codon model for positive selection.

### The evaluated intrinsic-fitness pipeline

#### Intrinsic fitness from a protein language model

We quantify intrinsic (replicative) fitness by the log-likelihood ratio that is assigned to a substitution by the PLM, using the masked-marginal scheme [4]. For a single substitution of the wild-type residue 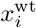 at position *i* to amino acid *a*, the intrinsic fitness term is defined as:

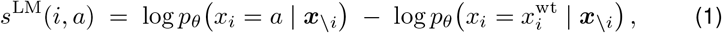

where *p* _*θ*_ is the model’s categorical distribution over the amino-acid alphabet at the masked position, and ***x***_*\i*_ denotes the sequence with position *i* replaced by the mask token. Because the score is a ratio of likelihoods for the mutant versus the wild-type residue in the same context, it measures how surprising a substitution is relative to what the model has learned. Because a PLM is trained on tens of millions of proteins, it has in effect internalized the protein family- and alignment-level constraints that an explicit evolutionary model would otherwise estimate, so even this single-sequence term carries some alignment-like information. When more than one substitution is present, that is when we have a multi-substitution genotype *M*, we sum the per-position contributions, *s*^LM^(*M* ) =Σ_(*i,a*) *M*_ *s*^LM^(*i, a*). Note that such an additive approximation neglects higher-order epistasis, at least beyond the one that is already captured by the model’s context.

By default, the PML we use is based on ESM-2 [6], but we also implement two other PMLs: the alignment-conditioned multiple-sequence-alignment (MSA) Transformer [16], and the autoregressive, retrieval-augmented Tranception model [17], which tolerates insertions and deletions and performs strongly on viral assays in the ProteinGym benchmark [7]. This lets us gauge how each model performs and, if useful, combine them. When multiple models are used, their per-mutation scores are standardized (transformed to zero mean and unit variance across *M* ) and averaged. This ensembling strategy is known to improve zero-shot accuracy [7]. This term is conceptually continuous with earlier unsupervised fitness models built on evolutionary couplings [12], latent-variable models [13, 14], and global epistatic models [15].

#### Benchmarking against deep mutational scanning

We then benchmark every estimator in the same way against a DMS score. This score reflects the experimental fitness, binding, or expression value that Prote-inGym [7] reports for a given single-amino-acid mutant, with higher scores denoting fitter or higher-affinity variants. We take the ProteinGym scores that released “as is,” pair each measured mutant with its predicted score by mutant identifier, and report per-assay accuracy as the Spearman rank correlation between the predicted and measured scores over all matched mutants.

#### Alignment-based intrinsic term: the PSSM

The alignment-based estimator of the intrinsic term is based on a site-independent log-odds model. For a substitution of the wild-type residue 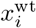 at position *i* to amino acid *a*, its score is

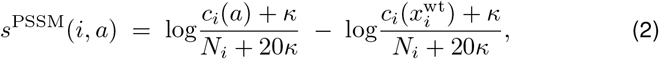

where *c*_*i*_(*b*) is the number of times amino acid *b* occurs in the alignment column mapped to position *i, N*_*i*_ =Σ_*b*_ *c*_*i*_(*b*) is that column’s occupancy, *κ* = 0.5 is a pseudocount added to each of the 20 amino-acid counts (hence 20*κ* in the denominator), and a multi-substitution genotype sums the per-position terms (positions with no aligned column contribute zero). The shared denominator cancels, so the score reduces to the smoothed log-ratio of the mutant to the wild-type residue frequency in that column.

#### Epoch-resolved viral alignments

To gauge the impact of heterotachy on fitness prediction, we assembled dated sequence alignments for two viruses with contrasting modes of evolution: SARS-CoV-2 spike and influenza A H3N2 hemagglutinin. For each, and evaluate how the collection date of the sequences used to build the PSSM affects prediction of a fixed DMS assay.

Spike protein sequences (1250-1275 residues, human host) were retrieved from NCBI via Entrez, sampling ∼200 sequences from each of fourteen date windows spanning 2020-2024, with each sequence’s collection date read from its GenPept source annotation (2,800 dated sequences across 31 distinct collection months). The receptor-binding domain (RBD; spike positions 331-531, the region assayed by Starr et al. [1]) was extracted from each isolate by local alignment to the Wuhan-Hu-1 RBD. As the RBD is free of indels across variants, this yields a column-aligned RBD for every sequence. For influenza the same procedure was applied to H3N2 hemagglutinin (HA): 2,730 dated HA protein sequences retrieved from NCBI (collection year taken from the GenPept qualifier or the strain name, spanning 1968-2024), the HA extracted by alignment to the assay reference.

For a given subset of sequences we built a site-independent PSSM (Eq 2), and scored it against the fixed DMS: the Starr et al. ACE2-binding assay (`SPIKE SARS2 Starr bind 2020`) for spike, and the Lee et al. [2] A/Perth/16/2009 replication assay (`C6KNH7 9INFA Lee 2018`, HA positions 1-566) for HA. We then swept the collection date under two schemes: a cumulative window (all sequences collected on or before date *T* ), and a sliding window (sequences within *±*6 months of *T* ). As above, we report Spearman correlation coefficients, this time between the PSSM score and the DMS as a function of *T* .

#### Phylogenetic estimators from a codon model

Both the intrinsic and the antigenic terms can also be estimated from a codon model that, relative to its sequence-count counterpart, separates selection from the neutral mutational process, while correcting for shared ancestry. Using the same dated SARS-CoV-2 and influenza datasets, the coding nucleotide sequences of every isolate were retrieved from its GenPept `coded by` annotation, in-frame codons were threaded onto a reference-guided protein alignment (columns thus correspond to reference-protein positions). Sequences with internal stop codons were removed, and a maximum-likelihood tree was estimated under a general time-reversible nucleotide-substitution model with empirical base frequencies and four-category gamma-distributed rate variation (GTR+F+Γ_4_) with IQ-TREE 2 (v2.3.6) [38].

For the intrinsic term, we fit the site-heterogeneous mutation-selection model of Rodrigue et al. [30] in PhyloBayes-MPI [39] (compiled from source, https://github.com/bayesiancook/pbmpi), with options `-mutsel -dp`, running two MCMC chains with a fixed topology, assessing convergence by the between-chain discrepancy and the effective sample size of the trace summary statistics (S3 Fig, S4 Fig). The posterior-mean site-specific amino-acid fitness profiles *π*_*i*_(*a*) give a per-mutation score Δ*F*_*i*_(wt → mut) = log π_*i*_(mut) − log π_*i*_(wt), the tree-based counterpart of the PSSM, which we score against the same DMS.

For the antigenic term we ran HyPhy (v2.5) [40] site-selection methods: SLAC, FEL [32], FUBAR, and the episodic-diversifying-selection test MEME [31]. We used the sites detected to be under positive selection to define an escape-propensity weight in [0, 1], either using a structure-based accessibility term (additive mode), or a multiplicative term (the multiplicative mode). Both estimators use the same codon alignment and tree, and the scripts that build these inputs, run each tool on all available cores, and parse the outputs are provided with the pipeline.

#### Implementation and availability

The method is implemented in Python (NumPy, pandas, and Biopython; PyTorch and the `transformers` library for PLM inference) as a modular pipeline with command-line entry points for scoring, structure-based RSA estimation, and growth-rate fitting. When PLM dependencies or weights are absent, the scorer falls back to a substitution-matrix baseline so that the downstream fusion and validation logic remain testable. Source code, worked examples, and documentation are provided in the accompanying repository (https://github.com/sarisbro), including fixed-seed code that regenerates the implementation-validation checks and the ProteinGym benchmark producing the empirical results (Table 1). The analyses used Python 3 with NumPy 2.2, pandas 2.3, SciPy 1.15, Biopython 1.87, and Matplotlib 3.10; language-model scoring used PyTorch and the HuggingFace `transformers` library, with the ESM-2 650M checkpoint `facebook/esm2 t33 650M UR50D`, and the MSA Transformer and Tranception values taken from ProteinGym’s released per-assay scores. The codon-model analyses used the external programs IQ-TREE 2 (v2.3.6) [38], HyPhy (v2.5) [40], and PhyloBayes-MPI (compiled from source). Exact package versions are pinned in the repository.

### The broader scoring framework (not evaluated here)

The components in this part complete the modular pipeline but are not scored against the real DMS benchmarks in this paper. Each is either checked on synthetic data (S1 Text) or left for future validation; we include them so that the intrinsic-term results can be read as one piece of a complete, reproducible framework rather than a stand-alone tool.

#### Framework logic

The method follows the modular, multiplicative logic of EVEscape [19], in which a mutation’s escape potential is the product of independent unsupervised factors, but replaces the protein-specific variational fitness model with the zero-shot likelihood of a protein language model [4, 6]. For a wild-type sequence ***x***^wt^ and a candidate substitution set *M* we compute three per-mutation terms (schematic in S1 Fig): an intrinsic fitness term derived from the PLM (above), an antibody accessibility term derived from structure, and a biochemical dissimilarity term; the last two are two separate views of antigenicity. These three terms are then combined into a single score under one of three fusion modes, and the fusion can optionally be oriented against an external, self-supervised measure of realized selection. All components are unsupervised in the sense that they require no DMS data; validation and calibration draw only on sequence data (natural alignments or genomic-surveillance frequencies).

#### Antigenic escape terms

We implement two structure- and chemistry-based terms that model antibody escape, taking inspiration from EVEscape [19]. Because a PLM likelihood penalizes rare substitutions, it assigns low intrinsic fitness to precisely the antigenically novel changes that are positively selected under immune pressure: the escape terms supply the complementary signal that the likelihood term cannot. A sequence-only alternative that plays the same role, the codon-model positive-selection flag, is described above (Phylogenetic estimators from a codon model) and can substitute for the structure-based accessibility term in the fusion.

##### 1. Accessibility (acc)

Antibodies predominantly engage solvent-exposed residues, so we weight substitutions by the relative solvent accessibility (RSA) of their position. Given a structure, we compute per-residue solvent-accessible surface area (SASA) with the Shrake-Rupley algorithm [41], and normalize by the residue-specific theoretical maximum [42],

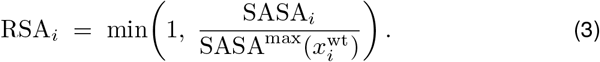

SASA is computed in the full multi-chain (biological-assembly) context by default, because accessibility is a property of the quaternary structure rather than of an isolated protomer. Since experimental accessibility is rarely available for the target, RSA is estimated from a homologous template: a candidate structure is retrieved from the Protein Data Bank [43] by MMseqs2-based sequence search [44], the template chain is aligned to the query, and per-residue RSA is transferred onto query positions through the alignment; positions without structural coverage revert to a neutral default. Structure parsing and SASA use Biopython [45], avoiding any external secondary-structure dependency [46].

##### Biochemical dissimilarity (dis)

Substitutions that change residue chemistry most strongly are those most likely to disrupt antibody binding. For a substitution 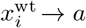 we define

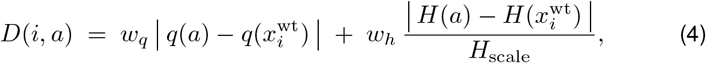

where *q*(·) is the formal charge at physiological pH, *H*(·) is the Kyte-Doolittle hydropathy [47], *H*_scale_ is the hydropathy range used as a normalizer, and *w*_*q*_, *w*_*h*_ are non-negative weights (see Sign-of-selection calibration below).

#### Score fusion

Let *t*_*k*_(*M* ) denote the three per-mutation terms (*k* ∈ {LM, acc, dis}) and *z*(·) the standardization operator applied across the scored mutation set. We define three fusion modes. The intrinsic mode returns the PLM term alone and targets replicative fitness. The additive mode forms a weighted sum in standardized space,

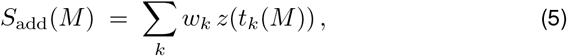

a decomposition of total fitness into intrinsic and antigenic contributions in the spirit of fitness models for influenza and SARS-CoV-2 [18]. The multiplicative mode reproduces the EVEscape product [19] by mapping each standardized term to a pseudo-probability through a temperature-scaled logistic function *σ*, summing in log space,

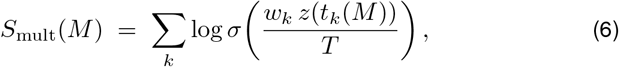

with temperature *T >* 0. The multiplicative mode enforces that a mutation scores highly only when it is simultaneously fitness-tolerated, accessible, and biochemically disruptive. The temperature *T* controls only how sharply each standardized term is squashed toward {0, 1}; it is a fixed hyperparameter (default *T* = 1) rather than a quantity fit to data. Because the empirical results reported here evaluate the intrinsic term alone (no fusion), *T* does not affect them; when the multiplicative mode is used in practice, *T* is either left at its default or chosen, together with the weights {*w*_*k*_}, on the same self-supervised target as the sign-of-selection calibration (below), never on held-out DMS labels.

#### Sign-of-selection calibration

Because the intrinsic term may not rank antigenically novel substitutions correctly, we optionally calibrate the fusion weights {*w*_*k*_} against an external signal of realized selection. Given target values *y*_*j*_ for a subset of mutations, we solve the ordinary least-squares problem

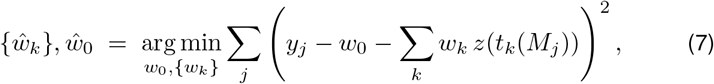

which both weights and orients (that is, fixes the sign of) each term. Targets are drawn, in order of preference, from surveillance-derived growth advantages (below), DMS measurements where available, or, as a fully sequence-only fallback, the log natural frequency of each substitution in an alignment. This keeps calibration self-supervised: no experimental phenotype is required.

### Growth advantages from genomic surveillance

This realized-fitness term is validated only against a simulated ground truth (S1 Text); its application to real surveillance data is left for future work. To obtain a realized-fitness signal for validation and calibration without experiments, we estimate per-mutation growth advantages from variant-count time series using a multinomial logistic regression (MLR) model, the standard approach in genomic-surveillance fitness estimation [35, 36, 48]. Let *n*_*it*_ be the observed count of sequence variant *i* at time *t*; the model assumes the variant frequencies follow

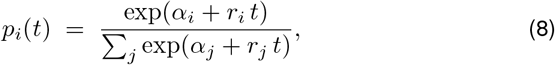

where *α*_*i*_ is a nuisance intercept and *r*_*i*_ is the log growth advantage of variant *i* per unit time. Following the per-mutation formulation of PyR0 [36], we optionally decompose the variant growth rate into additive contributions of the mutations it carries, *r*_*i*_ =Σ_*m*_ *G*_*im*_ *b*_*m*_, where *G*_*im*_ ∈ {0, 1} indicates whether variant *i* carries mutation *m*, and *b*_*m*_ is that mutation’s growth effect.

Parameters are fit by maximizing the multinomial log-likelihood with an *ℓ*_2_ penalty on the mutation effects,

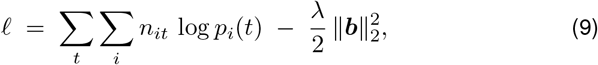

whose gradient with respect to the logits is *N*_*t*_ *p*_*i*_(*t*) *n*_*it*_ with *N*_*t*_ = Σ_*i*_ *n*_*it*_. Optimization uses Adam [49] on the exact gradient. Time is standardized for numerical conditioning and the growth rates are rescaled to the original time unit afterward. Only relative rates are identifiable under the softmax (Eq 8), so one variant is fixed as the reference and the ridge penalty resolves the residual null space of the mutation decomposition. The resulting {*b*_*m*_} provide the per-mutation growth targets used in Eq 7. As with all such phenomenological estimates, *r*_*i*_ and *b*_*m*_ conflate intrinsic transmissibility, immune escape, and epidemiological context, and assume an approximately constant advantage over the fitting window [35].

#### Label-free validation

We validate scores at three levels of stringency, none of which requires DMS. First, using only a multiple-sequence alignment of the target protein, we ask whether the score ranks substitutions observed in nature above those never seen. Writing *c*_*i*_(*a*) for the count of amino acid *a* at the aligned column corresponding to query position *i*, the natural frequency is *f*_*i*_(*a*) =*c*_*i*_(*a*)*/* Σ_*b*_ *c*_*i*_(*b*), and a substitution is deemed observed when its count and frequency exceed thresholds *c*_min_ and *f*_min_. By default *c*_min_ = 1 and *f*_min_ = 0 (any residue seen at least once in the aligned column counts as observed). Both are exposed as parameters so the definition of “observed” can be tightened (for example to exclude singletons or sequencing-error residues) on very deep alignments. We report the area under the receiver-operating-characteristic curve (AUROC) for observed (true positive) versus unobserved (false positive) substitutions, the Spearman correlation between score and log *f*_*i*_(*a*) over observed substitutions, and the precision among the top-*P* ranked predictions, where *P* is the number of observed substitutions. For calibration, log-frequency labels use a pseudocount *κ ŷ*_*i*_(*a*) = log[(c_*i*_(*a*) + *κ*) / (Σ_*b*_ c_*i*_(*b*) + 20*κ*)], 20*κ*), so that covered-but-unobserved substitutions receive a finite value. Second, where a phylogenetic estimate of empirical fitness is available, for example the observed-versus-expected mutation counts of Bloom and Neher [37], we report the Spearman correlation of the score against it. Third, we compare against realized epidemiological fitness by correlating scores with the surveillance-derived growth advantages of Eq 9 or with independent estimates of clade success [35, 50]. Disagreement across levels is informative: high concordance with alignment frequency but poor correlation with growth typically indicates a missing or mis-weighted antigenic term, consistent with the observation that evolutionary language captures replicative constraint while antigenic novelty requires an explicit escape signal [20].

## Data availability

All data underlying this study are publicly available. The deep mutational scanning assays and per-assay reference scores are from ProteinGym [7] (https://proteingym.org). The dated SARS-CoV-2 spike and influenza A H3N2 hemagglutinin sequences were retrieved from NCBI; the exact accession lists, retrieval queries, and dates, together with the derived alignments, trees, and per-mutation score tables, are deposited at https://github.com/sarisbro. All scripts were initially written by vibe-coding with Opus 4.8 [34], which was also used to proof this work. No new experimental data were generated, and the study used only publicly available sequence and assay data, so no ethics approval was required.

## Author contributions

Conceptualization: S.A.-B. Methodology: S.A.-B. and M.V. Software: S.A.-B. Formal analysis: S.A.-B. and M.V. Investigation: S.A.-B. and M.V. Writing, original draft: S.A.-B. Writing, review and editing: S.A.-B. and M.V. All authors read and approved the final manuscript.

## References

[1] Starr TN, Greaney AJ, Hilton SK, Ellis D, Crawford KHD, Dingens AS, et al. Deep Mutational Scanning of SARS-CoV-2 Receptor Binding Domain Reveals Constraints on Folding and ACE2 Binding. Cell. 2020;182(5):1295–1310.e20.

[2] Lee JM, Huddleston J, Doud MB, Hooper KA, Wu NC, Bedford T, et al. Deep mutational scanning of hemagglutinin helps predict evolutionary fates of human H3N2 influenza variants. Proceedings of the National Academy of Sciences. 2018;115(35):E8276–E8285.

[3] Greaney AJ, Loes AN, Crawford KHD, Starr TN, Malone KD, Chu HY, et al. Comprehensive mapping of mutations in the SARS-CoV-2 receptor-binding domain that affect recognition by polyclonal human plasma antibodies. Cell Host & Microbe. 2021;29(3):463–476.

[4] Meier J, Rao R, Verkuil R, Liu J, Sercu T, Rives A. Language models enable zero-shot prediction of the effects of mutations on protein function. In: Advances in Neural Information Processing Systems. vol. 34; 2021.

[5] Rives A, Meier J, Sercu T, Goyal S, Lin Z, Liu J, et al. Biological structure and function emerge from scaling unsupervised learning to 250 million protein sequences. Proceedings of the National Academy of Sciences. 2021;118(15):e2016239118.

[6] Lin Z, Akin H, Rao R, Hie B, Zhu Z, Lu W, et al. Evolutionary-scale prediction of atomic-level protein structure with a language model. Science. 2023;379(6637):1123–1130.

[7] Notin P, Kollasch AW, Ritter D, van Niekerk L, Paul S, Spinner H, et al. ProteinGym: Large-Scale Benchmarks for Protein Fitness Prediction and Design. In: Advances in Neural Information Processing Systems (Datasets and Benchmarks Track). vol. 36; 2023. .

[8] Brandes N, Goldman G, Wang CH, Ye CJ, Ntranos V. Genome-wide prediction of disease variant effects with a deep protein language model. Nature Genetics. 2023;55(9):1512–1522.

[9] Cheng J, Novati G, Pan J, Bycroft C, Avsec Ž, et al. Accurate proteome-wide missense variant effect prediction with AlphaMissense. Science. 2023;381(6664):eadg7492.

[10] Livesey BJ, Marsh JA. Using deep mutational scanning to benchmark variant effect predictors and identify disease mutations. Molecular Systems Biology. 2020;16(7):e9380.

[11] Starr TN, Greaney AJ, Stewart CM, Walls AC, Hannon WW, Veesler D, et al. Shifting mutational constraints in the SARS-CoV-2 receptor-binding domain during viral evolution. Science. 2022;377(6604):420–424.

[12] Hopf TA, Ingraham JB, Poelwijk FJ, Schärfe CPI, Springer M, Sander C, et al. Mutation effects predicted from sequence co-variation. Nature Biotechnology. 2017;35(2):128–135.

[13] Riesselman AJ, Ingraham JB, Marks DS. Deep generative models of genetic variation capture the effects of mutations. Nature Methods. 2018;15(10):816– 822.

[14] Frazer J, Notin P, Dias M, Gomez A, Min JK, Brock K, et al. Disease variant prediction with deep generative models of evolutionary data. Nature. 2021;599(7883):91–95.

[15] Laine E, Karami Y, Carbone A. GEMME: A Simple and Fast Global Epistatic Model Predicting Mutational Effects. Molecular Biology and Evolution. 2019;36(11):2604–2619.

[16] Rao RM, Liu J, Verkuil R, Meier J, Canny J, Abbeel P, et al. MSA Transformer. In: Proceedings of the 38th International Conference on Machine Learning (ICML). vol. 139 of PMLR; 2021. p. 8844–8856.

[17] Notin P, Dias M, Frazer J, Marchena-Hurtado J, Gomez AN, Marks DS, et al. Tranception: Protein Fitness Prediction with Autoregressive Transformers and Inference-time Retrieval. In: Proceedings of the 39th International Conference on Machine Learning (ICML). vol. 162 of PMLR; 2022. p. 16990–17017.

[18] fluksza M, Lässig M. A predictive fitness model for influenza. Nature. 2014;507(7490):57–61.

[19] Thadani NN, Gurev S, Notin P, Youssef N, Rollins NJ, Ritter D, et al. Learning from prepandemic data to forecast viral escape. Nature. 2023;622(7984):818– 825.

[20] Hie B, Zhong ED, Berger B, Bryson B. Learning the language of viral evolution and escape. Science. 2021;371(6526):284–288.

[21] Harvey WT, Carabelli AM, Jackson B, Gupta RK, Thomson EC, Harrison EM, et al. SARS-CoV-2 variants, spike mutations and immune escape. Nature Reviews Microbiology. 2021;19(7):409–424.

[22] Lopez P, Casane D, Philippe H. Heterotachy, an important process of protein evolution. Molecular Biology and Evolution. 2002;19(1):1–7.

[23] Galtier N, Gouy M. Inferring pattern and process: maximum-likelihood implementation of a nonhomogeneous model of DNA sequence evolution for phylogenetic analysis. Molecular Biology and Evolution. 1998;15(7):871–879.

[24] Foster PG. Modeling compositional heterogeneity. Systematic Biology. 2004;53(3):485–495.

[25] Blanquart S, Lartillot N. A Bayesian compound stochastic process for modeling nonstationary and nonhomogeneous sequence evolution. Molecular Biology and Evolution. 2006;23(11):2058–2071.

[26] Tuffley C, Steel M. Modeling the covarion hypothesis of nucleotide substitution. Mathematical Biosciences. 1998;147(1):63–91.

[27] Galtier N. Maximum-likelihood phylogenetic analysis under a covarion-like model. Molecular Biology and Evolution. 2001;18(5):866–873.

[28] Lartillot N, Philippe H. A Bayesian mixture model for across-site heterogeneities in the amino-acid replacement process. Molecular Biology and Evolution. 2004;21(6):1095–1109.

[29] Blanquart S, Lartillot N. A site- and time-heterogeneous model of amino acid replacement. Molecular Biology and Evolution. 2008;25(5):842–858.

[30] Rodrigue N, Philippe H, Lartillot N. Mutation-selection models of coding sequence evolution with site-heterogeneous amino acid fitness profiles. Proceedings of the National Academy of Sciences. 2010;107(10):4629–4634.

[31] Murrell B, Wertheim JO, Moola S, Weighill T, Scheffler K, Kosakovsky Pond SL. Detecting Individual Sites Subject to Episodic Diversifying Selection. PLOS Genetics. 2012;8(7):e1002764.

[32] Kosakovsky Pond SL, Frost SDW. Not So Different After All: A Comparison of Methods for Detecting Amino Acid Sites Under Selection. Molecular Biology and Evolution. 2005;22(5):1208–1222.

[33] Vilain M, Mghabghab R, Aris-Brosou S. Episodic positive selection accumulates through time in seasonal influenza A, with no climatic signature. bioRxiv. 2025;Preprint.

[34] Tomas M, Ndongo CPK, Vilain M, Aris-Brosou S. Variant emergence, not vaccine deployment, drives episodic positive selection on the SARS-CoV-2 spike at provincial scale in Canada. bioRxiv. 2026;Preprint.

[35] Abousamra E, Figgins M, Bedford T. Fitness models provide accurate shortterm forecasts of SARS-CoV-2 variant frequency. PLOS Computational Biology. 2024;20(9):e1012443.

[36] Obermeyer F, Jankowiak M, Barkas N, Schaffner SF, Pyle JD, Yurkovetskiy L, et al. Analysis of 6.4 million SARS-CoV-2 genomes identifies mutations associated with fitness. Science. 2022;376(6599):1327–1332.

[37] Bloom JD, Neher RA. Fitness effects of mutations to SARS-CoV-2 proteins. Virus Evolution. 2023;9(2):vead055.

[38] Minh BQ, Schmidt HA, Chernomor O, Schrempf D, Woodhams MD, von Haeseler A, et al. IQ-TREE 2: new models and efficient methods for phylogenetic inference in the genomic era. Molecular Biology and Evolution. 2020;37(5):1530–1534.

[39] Lartillot N, Rodrigue N, Stubbs D, Richer J. PhyloBayes MPI: Phylogenetic Reconstruction with Infinite Mixtures of Profiles in a Parallel Environment. Systematic Biology. 2013;62(4):611–615.

[40] Kosakovsky Pond SL, Poon AFY, Velazquez R, Weaver S, Hepler NL, Murrell B, et al. HyPhy 2.5—A Customizable Platform for Evolutionary Hypothesis Testing Using Phylogenies. Molecular Biology and Evolution. 2020;37(1):295– 299.

[41] Shrake A, Rupley JA. Environment and exposure to solvent of protein atoms. Lysozyme and insulin. Journal of Molecular Biology. 1973;79(2):351–371.

[42] Tien MZ, Meyer AG, Sydykova DK, Spielman SJ, Wilke CO. Maximum allowed solvent accessibilities of residues in proteins. PLOS ONE. 2013;8(11):e80635.

[43] Berman HM, Westbrook J, Feng Z, Gilliland G, Bhat TN, Weissig H, et al. The Protein Data Bank. Nucleic Acids Research. 2000;28(1):235–242.

[44] Steinegger M, Söding J. MMseqs2 enables sensitive protein sequence searching for the analysis of massive data sets. Nature Biotechnology. 2017;35(11):1026–1028.

[45] Cock PJA, Antao T, Chang JT, Chapman BA, Cox CJ, Dalke A, et al. Biopython: freely available Python tools for computational molecular biology and bioinformatics. Bioinformatics. 2009;25(11):1422–1423.

[46] Kabsch W, Sander C. Dictionary of protein secondary structure: pattern recognition of hydrogen-bonded and geometrical features. Biopolymers. 1983;22(12):2577–2637.

[47] Kyte J, Doolittle RF. A simple method for displaying the hydropathic character of a protein. Journal of Molecular Biology. 1982;157(1):105–132.

[48] Hadfield J, Megill C, Bell SM, Huddleston J, Potter B, Callender C, et al. Nextstrain: real-time tracking of pathogen evolution. Bioinformatics. 2018;34(23):4121–4123.

[49] Kingma DP, Ba J. Adam: A Method for Stochastic Optimization. In: 3rd International Conference on Learning Representations (ICLR); 2015. .

[50] Neher RA, Russell CA, Shraiman BI. Predicting evolution from the shape of genealogical trees. eLife. 2014;3:e03568.

