## Supplementary material for "A moving target: non-stationary selection governs unsupervised prediction of viral fitness": SI file

**S1 Text. Implementation validation on synthetic and simulated data.** These experiments characterize implementation correctness and the growth estimator against a known ground truth, not predictive accuracy on real proteins. All use fixed seeds and the substitution-matrix fallback scorer.

*End-to-end operation:* on a 60-residue test protein the pipeline enumerates and scores the complete single-substitution scan (1,140 mutants) across all three fusion modes. When protein-language-model weights are absent, it falls back to a substitution-matrix scorer without interrupting fusion, calibration, or validation, confirming that the modules are decoupled. Unit checks returned the analytically correct ranking-metric values (Spearman $1.0$; AUROC $1.0$).

*Growth estimator:* we simulated a variant-count time series from the multinomial-logistic growth model, in which the frequency of variant $i$ at time $t$ is

$$p_i(t) \;=\; \frac{\exp(\alpha_i + r_i\, t)}{\sum_j \exp(\alpha_j + r_j\, t)}, \tag{1}$$

with $\alpha_i$ a nuisance intercept and $r_i = \sum_m G_{im}\, b_m$ the log growth advantage of variant $i$ decomposed into additive per-mutation effects $b_m$. Parameters were fit by maximizing the $\ell_2$-penalized multinomial log-likelihood

$$\ell \;=\; \sum_t \sum_i n_{it}\, \log p_i(t) \;-\; \frac{\lambda}{2}\, \|\boldsymbol{b}\|_2^2, \tag{2}$$

where $n_{it}$ is the observed count of variant $i$ at time $t$. With prescribed additive per-mutation effects (twelve variants from seven identifiable mutations, twenty time points, 500 sequences each), the estimator recovered these at Spearman $0.93$ / Pearson $0.95$ per mutation and Spearman $0.95$ per variant (Table S5, S5 Fig). Feeding the fitted advantages $\{b_m\}$ into the self-supervised calibration

$$\{\hat{w}_k\}, \hat{w}_0 \;=\; \operatorname*{arg\,min}_{w_0, \{w_k\}} \sum_j \left( y_j - w_0 - \sum_k w_k\, z(t_k(M_j)) \right)^2, \tag{3}$$

placed positive weight on the fitness term.

*Structure and label-free validation:* the relative solvent accessibility of each position,

$$\mathrm{RSA}_i \;=\; \min\!\left(1,\; \frac{\mathrm{SASA}_i}{\mathrm{SASA}^{\max}(x_i^{\mathrm{wt}})}\right), \tag{4}$$

fell in $[0, 1]$ on a controlled structure, with the template-to-query mapping correct under a deliberate offset. On a synthetic alignment built to be consistent with the scorer, the label-free validator ranked observed substitutions above unobserved ones (AUROC $0.83$), a consistency check, not an unbiased performance estimate.

**Table S1. Description of the 21 viral ProteinGym assays.** Virus, protein, DMS assay identifier, and the phenotype each assay measures, from the ProteinGym reference metadata. All assays are scored so that higher values indicate fitter or higher-affinity variants. Nineteen of the 21 measure viral replication (organismal fitness), whereas the two SARS-CoV-2 spike RBD assays instead measure ACE2-binding affinity and RBD surface expression. Per-assay ESM-2 and PSSM scores are in Table S2.

| Virus | Protein | DMS assay identifier | Measured phenotype |
|---|---|---|---|
| SARS-CoV-2 | Spike RBD | SPIKE_SARS2_Starr_bind_2020 | ACE2-binding affinity |
| | Spike RBD | SPIKE_SARS2_Starr_expr_2020 | RBD surface expression |
| | Main protease (nsp5) | R1AB_SARS2_Flynn_growth_2022 | Replication (growth) |
| Influenza A | Hemagglutinin (H1, WSN/1933) | A0A2Z5U3Z0_9INFA_Doud_2016 | Replication |
| | Hemagglutinin (H1, WSN/1933) | A0A2Z5U3Z0_9INFA_Wu_2014 | Replication |
| | Hemagglutinin (H3, Perth/2009) | C6KNH7_9INFA_Lee_2018 | Replication |
| | Nucleoprotein (H3, Aichi/1968) | I6TAH8_I68A0_Doud_2015 | Replication |
| | Nucleoprotein (H1, PR8/1934) | NCAP_I34A1_Doud_2015 | Replication |
| | Neuraminidase (H1, 1933) | NRAM_I33A0_Jiang_standard_2016 | Replication |
| | Polymerase acidic PA (H1, PR8) | PA_I34A1_Wu_2015 | Replication |
| | Polymerase basic PB2 (H2, 1986) | A4D664_9INFA_Soh_CCL141_2019 | Replication (avian cells) |
| HIV-1 | Envelope (BF520) | A0A192B1T2_9HIV1_Haddox_2018 | Replication |
| | Envelope (BG505) | Q2N0S5_9HIV1_Haddox_2018 | Replication |
| | Envelope (89.6) | ENV_HV1B9_DuenasDecamp_2016 | Replication |
| | Envelope (BRU/LAI) | ENV_HV1BR_Haddox_2016 | Replication |
| | Rev | REV_HV1H2_Fernandes_2016 | Replication |
| | Tat | TAT_HV1BR_Fernandes_2016 | Replication |
| Zika virus | Envelope (E) | A0A140D2T1_ZIKV_Sourisseau_growth_2019 | Replication |
| Hepatitis C virus | NS5A (JFH-1) | POLG_HCVJF_Qi_2014 | Replication |
| Coxsackievirus B3 | Capsid polyprotein | POLG_CXB3N_Mattenberger_2021 | Replication |
| Adeno-assoc. virus 2 | Capsid (VP1) | CAPSD_AAV2S_Sinai_substitutions_2021 | Capsid production (viability) |

**Table S2. Per-assay ProteinGym benchmark.** ESM-2 versus the alignment PSSM on the 21 viral substitution assays, with the MSA quality-control columns (focus-to-target mapping identity and coverage). Each assay's virus, protein, and measured phenotype are described in Table S1.

| Assay | ESM-2 | PSSM | $\Delta$ | id. | cov. |
|---|---|---|---|---|---|
| *Viral assays* | | | | | |
| PA_I34A1_Wu_2015 | 0.04 | 0.62 | 0.58 | 1.000 | 1.000 |
| POLG_HCVJF_Qi_2014 | 0.14 | 0.64 | 0.51 | 1.000 | 1.000 |
| Q2N0S5_9HIV1_Haddox_2018 | 0.04 | 0.52 | 0.48 | 1.000 | 0.981 |
| A0A192B1T2_9HIV1_Haddox_2018 | 0.08 | 0.50 | 0.42 | 1.000 | 0.994 |
| NRAM_I33A0_Jiang_standard_2016 | 0.16 | 0.58 | 0.42 | 1.000 | 1.000 |
| ENV_HV1B9_DuenasDecamp_2016 | 0.01 | 0.36 | 0.35 | 1.000 | 1.000 |
| NCAP_I34A1_Doud_2015 | 0.03 | 0.38 | 0.35 | 1.000 | 1.000 |
| A0A140D2T1_ZIKV_Sourisseau_growth_2019 | 0.07 | 0.41 | 0.34 | 0.998 | 0.968 |
| I6TAH8_I68A0_Doud_2015 | 0.02 | 0.35 | 0.33 | 1.000 | 1.000 |
| TAT_HV1BR_Fernandes_2016 | 0.02 | 0.32 | 0.31 | 1.000 | 0.988 |
| ENV_HV1BR_Haddox_2016 | 0.05 | 0.34 | 0.29 | 1.000 | 0.978 |
| A4D664_9INFA_Soh_CCL141_2019 | 0.15 | 0.44 | 0.29 | 1.000 | 1.000 |
| SPIKE_SARS2_Starr_expr_2020 | 0.02 | 0.29 | 0.27 | 1.000 | 1.000 |
| SPIKE_SARS2_Starr_bind_2020 | -0.02 | 0.13 | 0.15 | 1.000 | 1.000 |
| CAPSD_AAV2S_Sinai_substitutions_2021 | 0.28 | 0.36 | 0.09 | 1.000 | 1.000 |
| POLG_CXB3N_Mattenberger_2021 | 0.36 | 0.42 | 0.06 | 0.998 | 0.971 |
| A0A2Z5U3Z0_9INFA_Wu_2014 | 0.46 | 0.50 | 0.04 | 1.000 | 0.987 |
| A0A2Z5U3Z0_9INFA_Doud_2016 | 0.51 | 0.50 | -0.01 | 1.000 | 0.975 |
| REV_HV1H2_Fernandes_2016 | 0.24 | 0.20 | -0.04 | 1.000 | 0.947 |
| R1AB_SARS2_Flynn_growth_2022 | 0.10 | 0.04 | -0.06 | 1.000 | 1.000 |
| C6KNH7_9INFA_Lee_2018 | 0.48 | 0.37 | -0.11 | 1.000 | 0.977 |
| *Viral mean* | 0.15 | 0.39 | 0.24 | | |
| *Non-viral control* | | | | | |
| BLAT_ECOLX_Firnberg_2014† | 0.74 | 0.49 | -0.24 | 1.000 | 0.817 |

Difference $\Delta = \text{PSSM} - \text{ESM-2}$; id. and cov. are the PSSM focus-to-target alignment identity and per-assay coverage. Viral assays are sorted by $\Delta$; the non-viral assay is a positive control. †: MSA identity or coverage $< 0.9$.

**Table S3. Codon-model site-selection counts (HyPhy).** Number of codons called under positive/diversifying selection by each HyPhy site test, on the full codon alignment of each virus (H3N2 HA, $566$ codons; SARS-CoV-2 spike, $1{,}273$ codons). Significance thresholds: $p < 0.1$ (SLAC, FEL, MEME) or posterior $\mathrm{Prob}[\beta > \alpha] > 0.9$ (FUBAR), each additionally requiring $\beta > \alpha$. The resulting per-site prior correlates only weakly with the mean per-site tolerance DMS (Spearman $\leq 0.10$ for both proteins).

| Site test | H3N2 HA | SARS-CoV-2 spike |
|---|---|---|
| SLAC | $5$ | $2$ |
| FEL | $6$ | $4$ |
| MEME | $35$ | $7$ |
| FUBAR | $4$ | $21$ |

**Table S4. Sites detected under positive or diversifying selection by each HyPhy method.** Reference-protein codon positions called significant at the thresholds of Table S3, for H3N2 HA ($566$ codons) and SARS-CoV-2 spike ($1{,}273$ codons); positions are in reference-protein coordinates (H3N2 HA numbering includes the signal peptide, so it is offset from mature-H3 antigenic-site numbering). Within each virus, positions detected by all four methods are in **bold** and those detected by exactly two methods are in italics; the remaining positions are called by one or three methods. On the spike, the detected sites are enriched in the receptor-binding domain (positions $331$-$531$), consistent with an antibody-escape interpretation; the sites supported by all four methods ($452$, $455$) both lie in this domain.

| Virus | Test | Reference-protein positions |
|---|---|---|
| H3N2 HA | SLAC | **127**, 151, **153**, **160**, *175* |
| | FEL | **127**, 151, **153**, **160**, *215*, 277 |
| | MEME | 2, 66, 69, 78, 107, 121, **127**, 147, 151, **153**, **160**, 171, 174, *175*, 208, 210, *215*, 236, 237, 241, 242, 274, 276, 277, 283, 286, 289, 320, 363, 380, 434, 534, 539, 545, 564 |
| | FUBAR | **127**, **153**, **160**, 277 |
| SARS-CoV-2 spike | SLAC | **452**, **455** |
| | FEL | 157, 445, **452**, **455** |
| | MEME | 69, 157, *213*, *444*, 445, **452**, **455** |
| | FUBAR | 83, 95, 146, 148, 157, 212, *213*, 222, 346, 417, *444*, 445, 446, **452**, **455**, 456, 478, 486, 655, 681, 701 |

**Table S5. Implementation-validation experiments on synthetic and simulated data.**

| Experiment | Setup | Result |
|---|---|---|
| Growth estimator (per mutation) | 12 variants, 7 effects, 20 time points | Spearman $0.93$, Pearson $0.95$ |
| Growth estimator (per variant) | same simulation | Spearman $0.95$ |
| Structure-based RSA | controlled structure + offset align | correct range and position mapping |
| Label-free MSA validation | synthetic alignment (consistency check) | AUROC $0.83$, enrichment@$P$ $0.48$ |
| Synthetic DMS scoring | baseline score $+$ noise | Spearman $0.58$ |
| Ranking-metric unit checks | monotonic / separable inputs | Spearman $1.0$, AUROC $1.0$ |

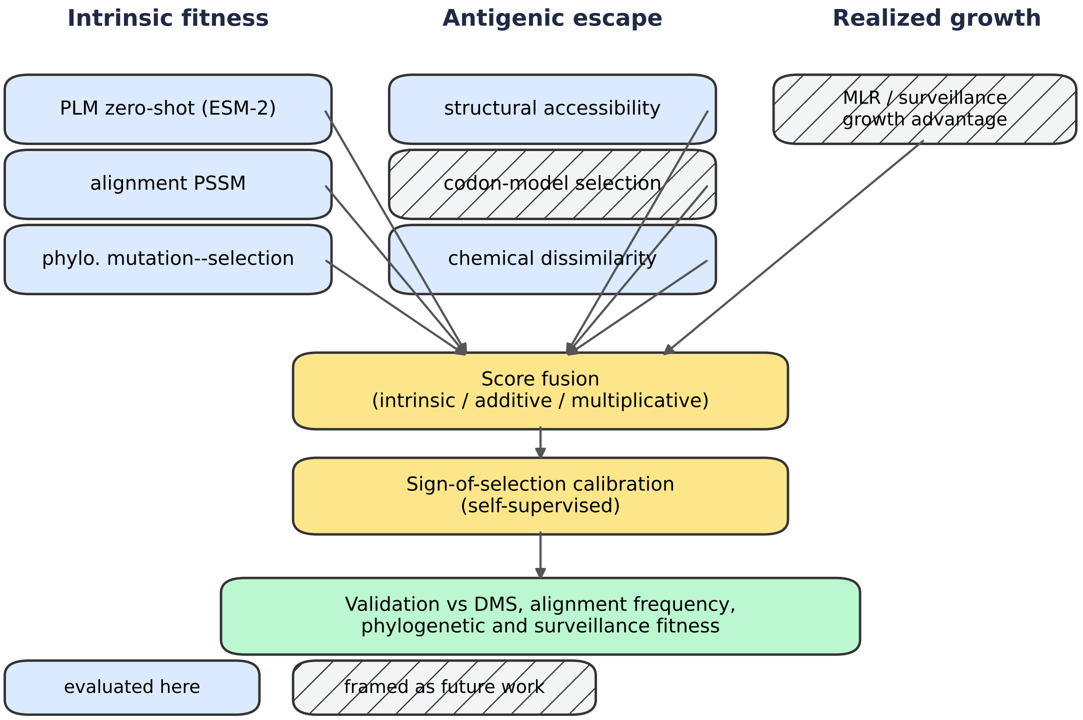


**S1 Fig. Overview of the unsupervised pipeline.** Each of three per-mutation terms (intrinsic replicative fitness, antigenic escape, and realized epidemiological growth) can be produced by more than one label-free estimator, so the estimator is treated as a swappable modeling choice. The terms are combined under one of three fusion modes, optionally oriented by a self-supervised sign-of-selection calibration, and validated against DMS, alignment frequency, and phylogenetic or surveillance fitness. Solid boxes are estimators evaluated in this work; hatched boxes (codon-model antigenic selection, surveillance growth) are components framed as future validation.

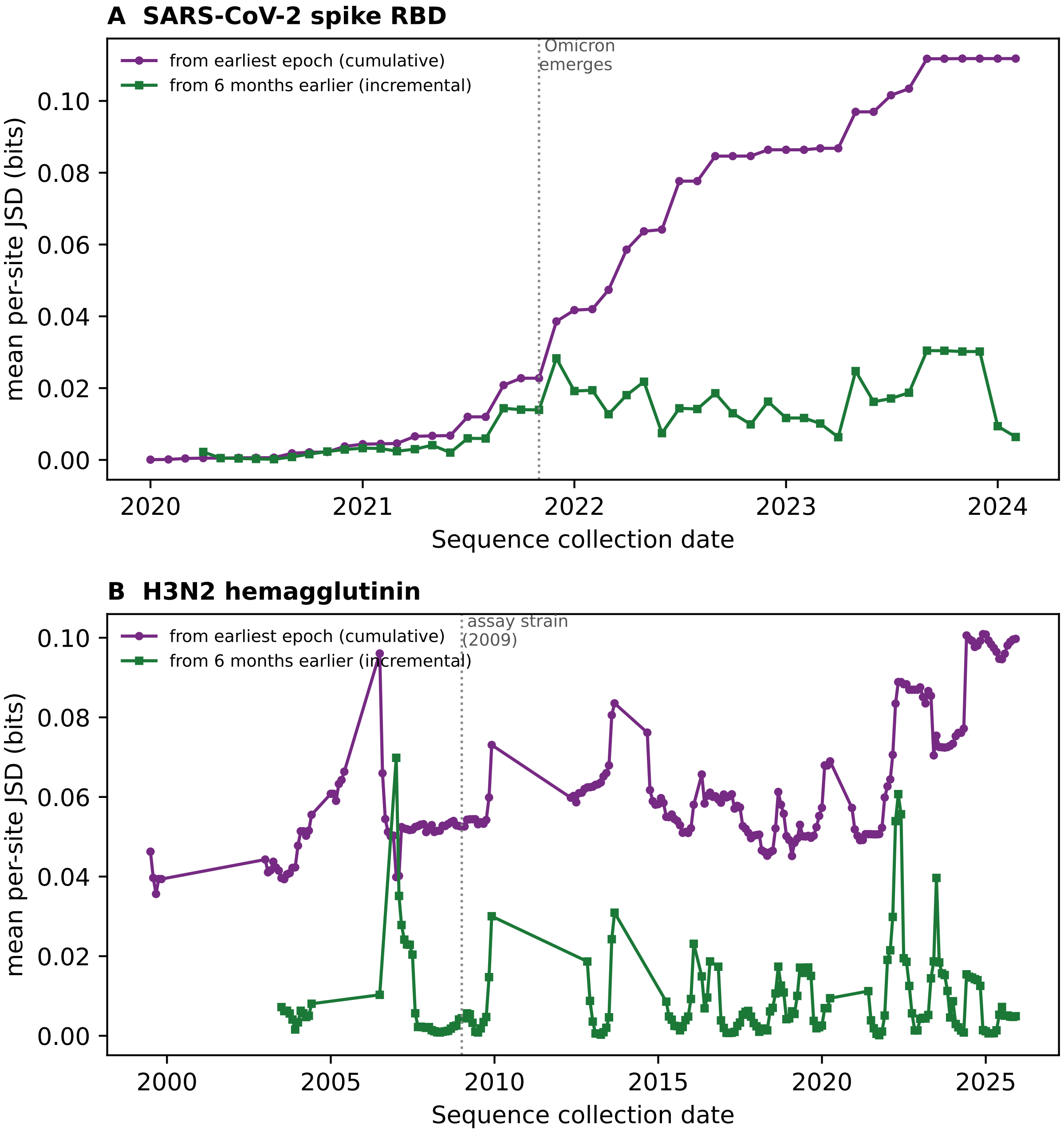


**S2 Fig. Non-stationarity measured as cumulative versus incremental drift.** For each virus, the mean per-site Jensen-Shannon divergence (bits) of each sliding $\pm 6$-month amino-acid profile from the earliest-epoch profile (cumulative, purple; as in the main-text non-stationarity figure) and from the profile six months earlier (incremental, green). **(A)** SARS-CoV-2 spike RBD: the incremental curve is near zero until a sharp transient at the emergence of Omicron. **(B)** H3N2 hemagglutinin: the incremental curve shows a recurring sawtooth as antigenic clusters turn over. The incremental measure recovers the same two modes as the cumulative one, showing that the earliest-epoch reference reports genuine accumulated drift rather than inflating the effect.

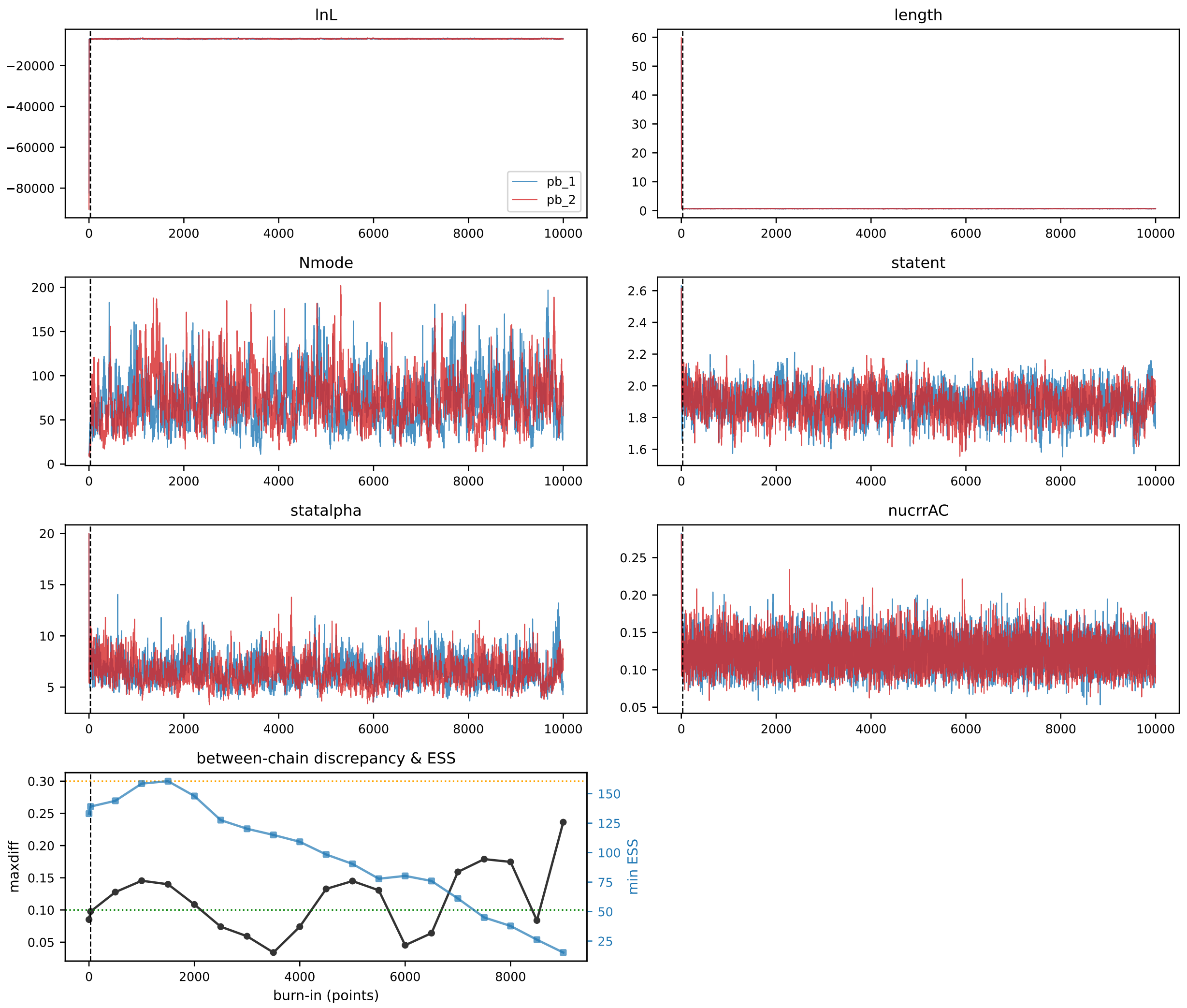


**S3 Fig. PhyloBayes mutation-selection convergence diagnostics, SARS-CoV-2 spike.** Trace plots of the log-likelihood and the model summary statistics (tree length, number of occupied mixture components, stationary entropy, the Dirichlet-process concentration, and a representative exchangeability) for the two independent MCMC chains ($10{,}000$ points each), with the selected burn-in marked (dashed line). Bottom panel: the `tracecomp`-style between-chain discrepancy $d = 2|\mu_1 - \mu_2|/(\sigma_1 + \sigma_2)$ (maximized over statistics) and the minimum effective sample size (Geyer's initial monotone sequence estimator) as a function of burn-in. The chains reach the stationary regime within $\sim 30$ points and reproduce one another ($\mathrm{maxdiff} < 0.1$).

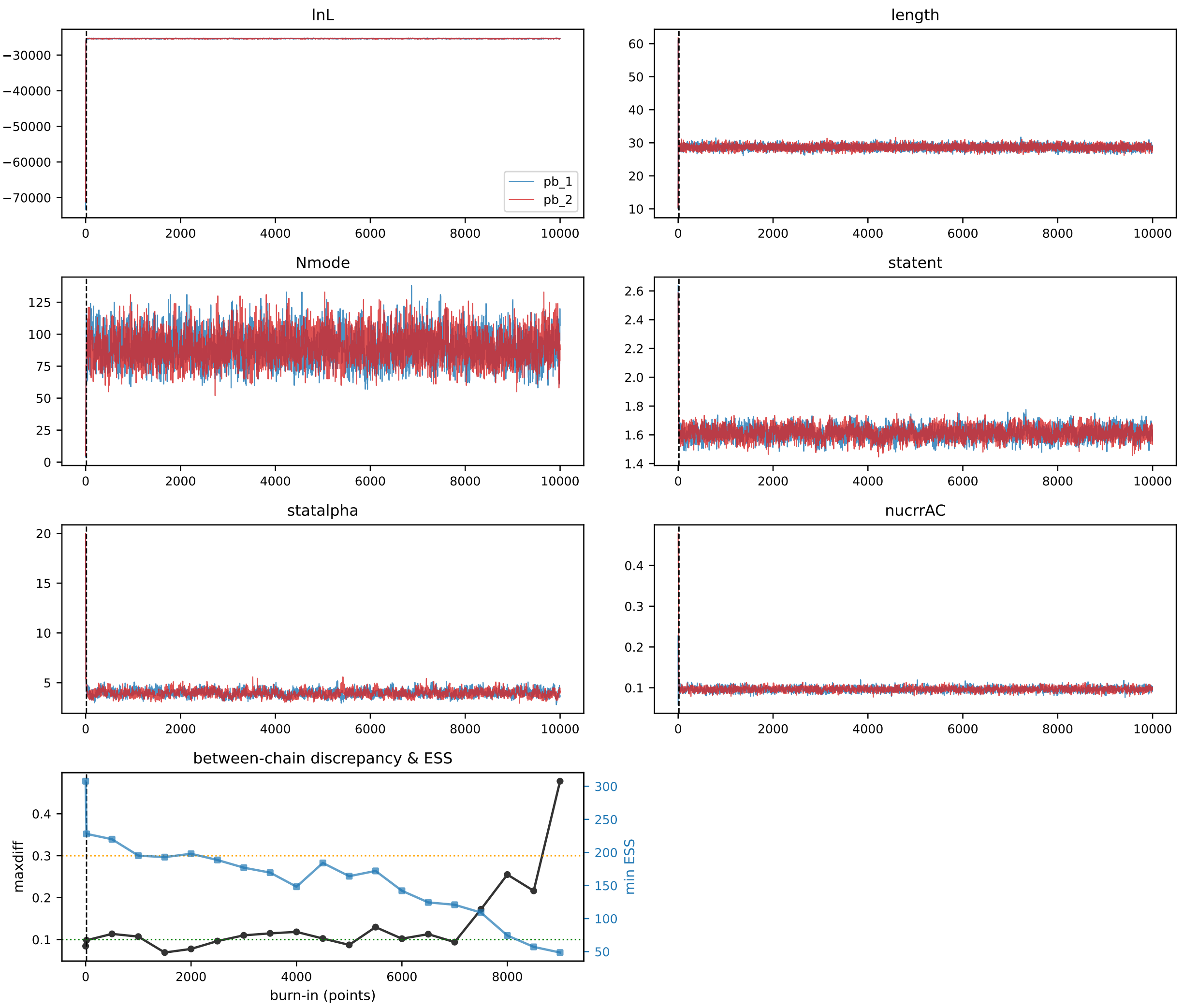


**S4 Fig. PhyloBayes mutation-selection convergence diagnostics, H3N2 hemagglutinin.** As S3 Fig, for the H3N2 HA chains ($10{,}000$ points each). The chains reach stationarity within $\sim 20$ points and agree across chains ($\mathrm{maxdiff} < 0.1$).

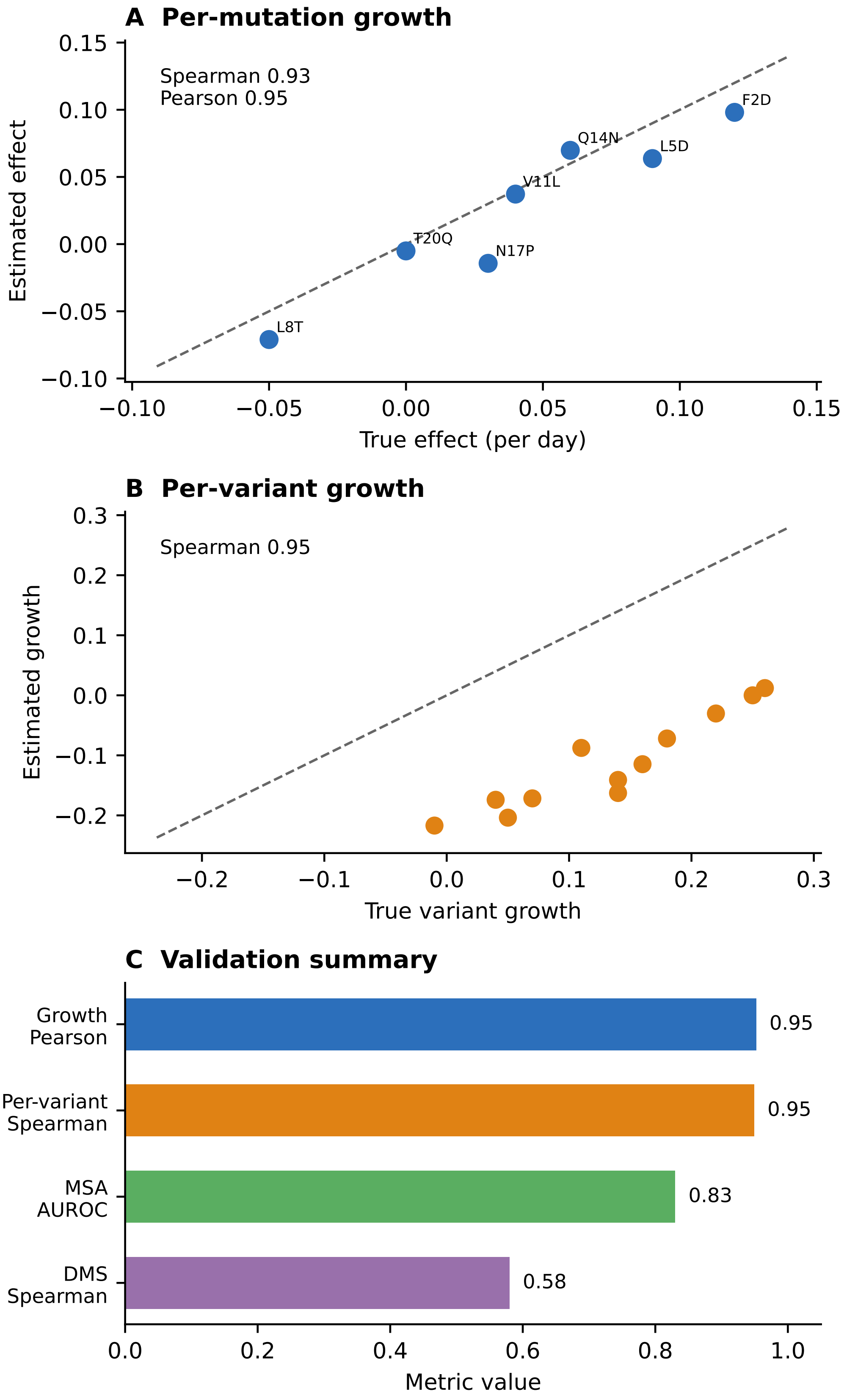


**S5 Fig. Implementation-validation results.** (A) Recovery of prescribed per-mutation growth effects by the multinomial-logistic estimator on simulated variant-count data (points, the seven identifiable mutations; dashed line $y = x$). (B) Recovery of the per-variant growth rates for the same simulation. (C) Summary of the validation metrics of Table S5.
